# A Human Hippocampal Organoid Model with Sustained Neural Stem Cells Reveals State Shifts Under Glucocorticoid Stress

**DOI:** 10.1101/2025.11.03.686207

**Authors:** Olivia Soper, Muhammad Z.K. Assir, Delia Ramírez, Gabriella Crawford, Richard Gyuris, Sara S.M. Valkila, Laetitia L. Lecante, Peng Liu, Paul A. Fowler, J. Antonio González, Daniel A. Berg, Eunchai Kang

## Abstract

The human hippocampus is a critical brain region for learning, memory, and stress regulation, distinguished by its ability to sustain neurogenesis after birth. This plasticity is driven by hippocampal neural stem cells (NSCs), which generate new neurons and maintain circuit integrity, but are highly sensitive to environmental and pathological influences. Mechanistic insight into human hippocampal development and neurogenesis remains limited by the absence of physiologically relevant models. Here, we establish an optimized protocol to generate human induced pluripotent stem cell-derived hippocampal organoids that recapitulate key features of hippocampal development. These organoids maintain organized NSC niches, support ongoing neurogenesis, and generate hippocampus-specific cell types. Cellular, transcriptomic, and electrophysiological analyses confirm progressive neuronal maturation, synapse formation, and functional activity, highlighting the physiological relevance of the system. Using this model, we modeled excess prenatal glucocorticoid exposure with dexamethasone, which perturbed NSC dynamics by reducing proliferation and inducing a precocious quiescent-like state. RNA sequencing revealed downregulation of NSC activation genes and upregulation of quiescence- and autophagy-associated programs, suggesting that glucocorticoid signaling enforces an early transition toward quiescence. These findings reveal a mechanism by which excessive glucocorticoid exposure may impair hippocampal growth. Together, this study introduces a robust human hippocampal organoid platform for dissecting the regulation of hippocampal development and for modeling the impact of environmental stressors on human neurogenesis.

## Introduction

The hippocampus is essential for mediating higher cognitive functions such as learning and memory, and for regulating emotion, fear, and stress. Unlike other brain regions, the hippocampus maintains a high level of plasticity via ongoing neurogenesis from neural stem cells (NSCs) after birth^1–3^. While this extended developmental plasticity supports circuit maturation and adaptability, it also renders the hippocampus especially sensitive to environmental influences and stress-related signalling^4,5^.

Dentate gyrus NSCs originate from dentate neuroepithelial progenitors, expand their population, and generate PROX1^+^ granule neurons and glial cells during development, before transitioning in early postnatal life into quiescent radial glia-like cells that retain the capacity for later reactivation^6^. As this NSC pool is established prenatally^7^, perturbations of the intrauterine environment that disrupt hippocampal development are increasingly recognized as determinants of later vulnerability. This idea aligns with the developmental origins of health and disease (DOHaD) hypothesis, which highlights the link between the prenatal environment and the risk of disease later in life^8^. Maternal influences are central to this framework, as the maternal environment directly shapes fetal development with long-lasting impacts on health outcomes throughout childhood and adulthood^9^. A wide range of environmental factors, including maternal nutrition, smoking, infection, and stress can contribute to altered developmental trajectories and increased susceptibility to disease^10^.

Glucocorticoid signaling is one of the key maternal influences on fetal brain development. Endogenous glucocorticoids readily cross the blood-brain barrier and play essential roles in normal brain development, particularly in the hippocampus, which expresses high levels of glucocorticoid receptors^11,12^. These hormones regulate critical neurodevelopmental processes, including neurogenesis, neuronal migration, neurotransmitter activity, and synaptic plasticity within the hippocampus^13,14^. However, excessive glucocorticoid signaling impairs hippocampal development. In humans, elevated prenatal stress has been associated with smaller fetal hippocampal volumes^15^, altered hippocampal functional connectivity in neonates^16^, and slower hippocampal growth during the first six months of life^17^. Furthermore, recent epidemiological studies suggest that anatomical alterations caused by maternal stress are associated with reduced cognitive function^18^ and an increased risk of neurodevelopmental disorders in the offspring^19^.These clinical observations have been investigated in animal models. In rodent and non-human primate models, elevated glucocorticoid levels due to prenatal stress have been linked to adverse effects on the hippocampus. Specifically, reductions in total hippocampal volume and impaired postnatal neurogenesis in the dentate gyrus have been reported following prenatal stress^14,20,21^. In addition, long-term structural alterations, including decreased synaptic density and reduced dendritic branching in the CA1 and CA3 regions, have been observed in the hippocampus of prenatally stressed rats^22–24^. Elucidating the cellular and molecular mechanisms by which excessive glucocorticoid signaling disrupts hippocampal development could deepen our understanding and guide therapeutic strategies for related pathologies. However, due to the lack of suitable human hippocampal models, much of our current knowledge relies on rodent models, which differ markedly from humans in brain structure and neurogenic dynamics.

There have been pioneering attempts to generate human hippocampal neurons. Functional hippocampal granule and pyramidal neurons have been derived from human stem cells using 2D long-term cultures^25,26^, and CA3 neurons have been generated from stem cells to model hippocampal connectivity^27^. Recent advances have enabled the generation of 3D hippocampal organoids from human induced pluripotent stem cells (iPSCs), providing an initial framework for studying human hippocampal development^28^. Similarly, hippocampal spheroids containing PROX1^+^ and ZBTB20^+^ neurons have demonstrated successful specification of key hippocampal cell types^29^. More recently, hippocampal organoids integrated with a neural interface have enabled recording of neural activity from PROX1^+^ZBTB20^+^ neurons^30^. Current hippocampal differentiation protocols rely on similar patterning cues and generally progress through three phases: neural induction, hippocampal specification, and maturation. The specification phase involves sustained activation of WNT and BMP signaling to promote dorsal telencephalic and hippocampal identity^31^. While these protocols show encouraging potential, they still face critical limitations, including NSC depletion, disorganized NSC structures, limited capacity for long-term culture and lack of glial cells. Therefore, further refinement is needed to develop human hippocampal organoids that can more faithfully model the dynamic and continuous nature of hippocampal neurogenesis.

In this study, we developed and comprehensively characterized a new human iPSC-derived hippocampal organoid system. This model recapitulates key aspects of human hippocampal development, including sustained NSC maintenance, ongoing neurogenesis, generation of hippocampus-specific neuronal subtypes and astrocytes and maturation of functional neurons, confirmed at cellular, transcriptomic, and electrophysiological levels. These features provide a robust platform for dissecting the cellular and molecular mechanisms governing human hippocampal formation. Using this system, we show that excessive glucocorticoid exposure disrupts normal neurogenesis and drives NSCs toward a quiescent-like state. Together, our work establishes a versatile human hippocampal organoid model for investigating both physiological and pathological regulation of hippocampal development and neurogenesis.

## Results

### Optimization of human hippocampal organoid generation

Hippocampal regional identity is shaped by spatial and temporal gradients of morphogens such as WNTs and BMPs **(Figure 1A)**, which drive dorsal telencephalic specification and regulate progenitor proliferation, neuronal differentiation, and patterning. To optimize hippocampal organoid generation, we systematically refined existing protocols by modifying key morphogens, adjusting WNT and BMP exposure timelines, and introducing structural handling improvements using the iPSC line, KOLF2-1J. The initial protocol (P1), adapted from published methods^28,29,32^ but updated with more potent and widely used signaling modulators **(Figure S1A)**, resulted in disorganized, non-radial growth, characterized by peripheral localization of SOX2^+^ NSCs and inward migration of DCX^+^ newly generated neurons. This inverted polarity led to limited expansion, structural instability, and fragmentation over time, ultimately compromising long-term culture viability **(Figure S1E-L)**. In Protocol 2 (P2), WNT signaling inhibition was introduced during the neural induction phase via IWP2 to promote NSC layer expansion, while exposure to WNT and BMP signaling during the specification phase was extended to enhance hippocampal commitment **(Figure S1A)**. These modifications significantly improved NSC niche formation and organoid growth by establishing correct radial polarity **(Figure S1E-K)**. In Protocol 3 (P3), as an attempt to enhance the radial polarity formation, a dissociation reaggregation step was introduced at day 10 when most cells were still SOX2^+^ NSCs **(Figure S1A-D)**, while keeping all other steps identical to P2. This additional step further improved radial organization by facilitating structural reorganization of NSCs prior to neuronal differentiation. However, the growth capacity and ability to generate PROX1^+^ granule neurons were reduced, potentially due to thinner NSC layers within each rosette-like structure, as reflected by increased organoid circularity and decreased size of NSC layers **(Figure 1E-K and M)**. In P4, Matrigel embedding, which is routinely used in cerebral organoid culture to promote neuroepithelial expansion during the patterning phase (Lancaster and Knoblich, 2014), was introduced in P2 **(Figure S1N)**. Embedding in Matrigel transiently altered hippocampal organoid morphology but ultimately disrupted NSC organization and radial growth, suggesting limited benefit for hippocampal organoid cultures **(Figure S1O and P)**. Among the tested conditions, P2 consistently exhibited the most robust radial architecture, well-organized NSC layers, and efficient generation of PROX1^+^ hippocampal neurons. Therefore, we adopted it as the optimized protocol for all subsequent experiments.

**Figure 1.**
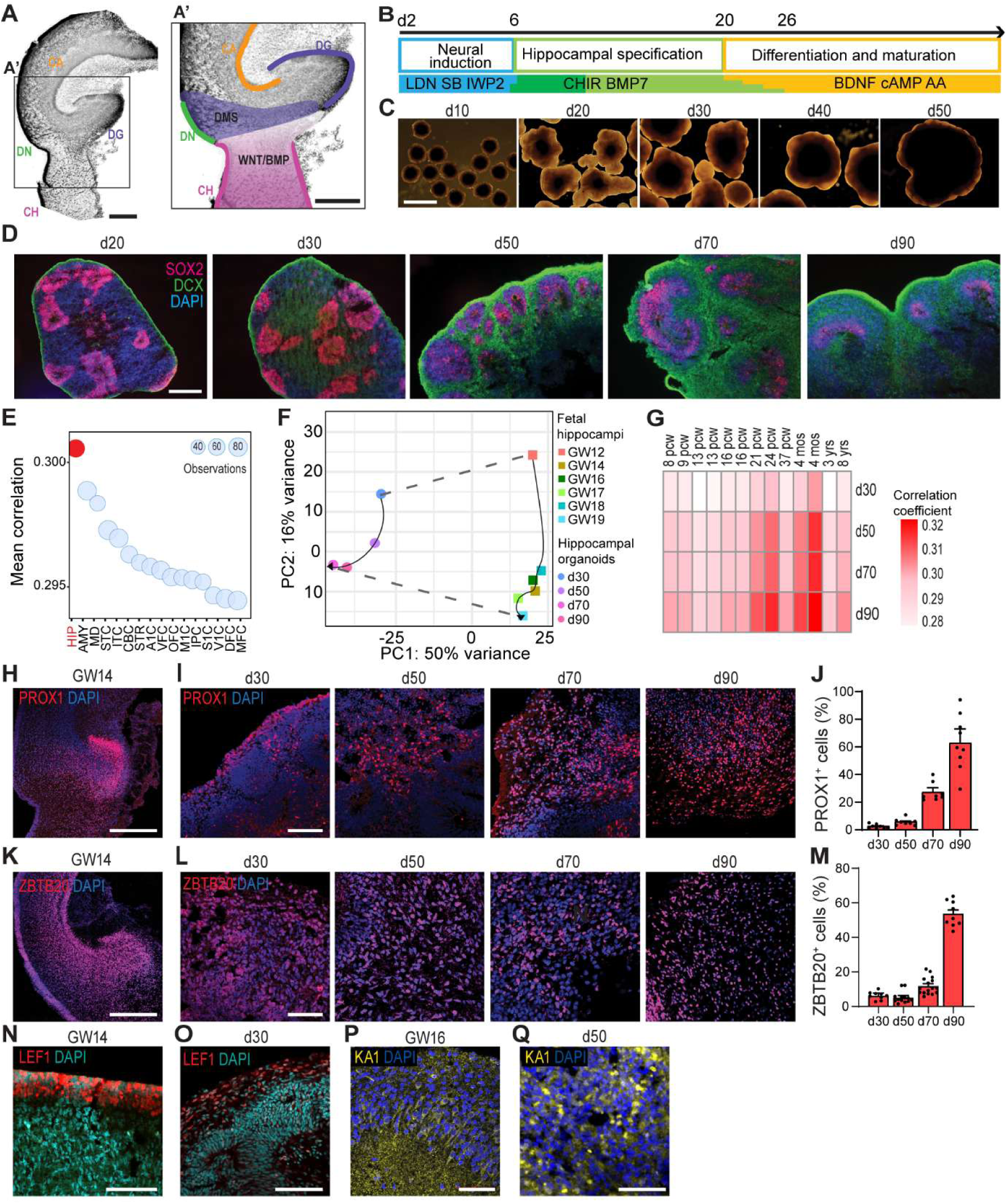
Generation and characterization of human hippocampal organoids. **(A)** Schematic of the developing human hippocampus at gestational week (GW) 16 highlighting key structures (**A**) and the importance of WNT/BMP signaling in cell patterning (**A’**). Scale bar = 500 μm. **(B)** Overview of the protocol for hippocampal organoid generation. Gradual transitions between media phases are indicated by stepwise color changes (see STAR Methods). **(C)** Representative brightfield images of hippocampal organoids at different developmental stages (d10-d50). Scale bar = 500 μm. **(D)** Representative immunofluorescence images showing NSCs (SOX2, magenta) and immature neurons (DCX, green) at d20-d90. Scale bar = 500 μm. **(E)** Bubble plot showing correlation of organoid transcriptomes with human brain regions from the Allen Brain Atlas. Only male samples were included to match the male iPSC line (KOLF2-1J) used for hippocampal organoid generation. Organoids exhibited the strongest correlation with the hippocampus (HIP, red) **(F)** PCA of bulk RNA-seq data from organoids (d30-d90) and fetal hippocampi (GW12-GW19). Organoids show progressive maturation parallelling the developmental trajectory of the fetal hippocampus. **(G)** Correlation heatmap of transcriptomics between organoids and human hippocampal tissue across lifespan from male samples, showing increasing hippocampal identity with organoid age. **(H-I)** Immunofluorescence images of dentate granule cells (PROX1, red) in GW14 hippocampus (**I**) and hippocampal organoids (d30-d90) (**H**). Scale bar = 500 μm (**H**), 100 μm (**I**) **(J)** Quantification of PROX1^+^ cells over time (d30-d90) shows significant increases (mean + SEM; n = 8-9 images from ≥3 organoids per timepoint). **(K and L)** Immunofluorescence images of hippocampal neurons (ZBTB20, red) in GW14 hippocampus (**K**) and hippocampal organoid (d30-d90) (**L**). Scale bar = 500 μm (**K**), 100 μm (**L**) **(M)** Quantification of ZBTB20^+^ cells over time (d30-d90) shows significant increases (mean + SEM; n = 9-16 images from ≥3 organoids per timepoint). **(N and O)** Representative Immunofluorescence images of LEF1^+^ cells (red) in GW14 hippocampus (**N**) and d30 hippocampal organoid (**O**). Scale bar = 100 μm. **(P and Q)** Representative Immunofluorescence image of CA pyramidal neurons (KA1, yellow) in GW 16 hippocampus (**P**) and d50 hippocampal organoid (**Q**). Scale bar = 50 μm.

To evaluate the effect of embryoid body (EB) size on radial growth, we generated EBs from 30,000, 60,000, and 120,000 iPSCs. Smaller EBs (30,000 cells) produced non-radial organoids, whereas 60,000- and 120,000-cell EBs supported proper radial organization, though 120,000 cell EBs showed reduced growth capacity at day 40 **(Figure S2A-D)**. EB area at day 2 emerged as a key predictor of polarity, with an optimal range of 0.37-0.54 mm² ensuring reproducible radial organization **(Figure S2E-F)**. Notably, P1 organoids failed to establish radial growth even within this optimal range (Data not shown). Together, these optimizations established a reliable protocol for generating human hippocampal organoids with correct radial polarity, and well-structured NSC niches.

### Validation of human hippocampal organoids

Following optimization, P2 was selected as the standard protocol, as it reliably generated hippocampal organoids with robust radial growth and well-established NSC layers **(Figure 1B-D)**. To validate hippocampal identity, we compared organoid transcriptomes from days 30-90 with reference datasets from the Allen Brain Atlas, which span 8 post-conception weeks (PCW) to 40 years. Correlation analyses revealed the strongest alignment between organoids and the hippocampus, although other regions such as the cerebellar cortex, striatum, and primary motor cortex also showed notable similarity, reflecting shared transcriptional features across brain regions as reported in previous study^33^ **(Figure 1E)**. To assess whether organoids recapitulate the developmental trajectory of the human fetal hippocampus, we compared transcriptomic data from organoids at days 30, 50, 70, and 90 with fetal hippocampi from gestational week (GW) 12-19. Principal component analysis revealed progressive maturation of the organoids, closely paralleling fetal hippocampal development, with remaining transcriptomic differences potentially attributable to organoid versus tissue origin due to the absence of microglia or vasculature **(Figure 1F)**.

Furthermore, correlation with hippocampal identity strengthened as organoids matured, peaking between 21 PCW and 4 months **(Figure 1G)**. These findings confirm that organoids progressively acquire and maintain hippocampal-like identity in vitro.

We next examined the emergence of hippocampal cell types using marker-based immunostaining. PROX1^+^ dentate granule cells and ZBTB20^+^ hippocampal neurons were observed in fetal tissue and progressively enriched in organoids from day 30 to 90, reaching ∼63% and ∼54% respectively **(Figure 1H-M)**. Additional hippocampal markers, including LEF1, enriched in dentate progenitors^34^; and KA1, a marker for CA3 pyramidal neurons^35^, were also detected **(Figure 1N-Q)**, supporting the presence of diverse hippocampal subtypes.

To test reproducibility, P2 organoids were also generated from two other iPSC lines, KOLF2-C1, and IMR90. Despite early line-to-line differences in growth, all produced organoids of comparable size and morphology by day 50 **(Figure S3A-B)**. Each line generated organoids with radial architecture, central SOX2^+^ NSCs, peripheral DCX^+^ immature neurons, and PROX1^+^ dentate granule cells **(Figure S3C-D)**. Thus, P2 robustly and reproducibly yields hippocampal organoids across multiple iPSC lines.

### Sustained NSC populations and ongoing neurogenesis in hippocampal organoids

A defining feature of the hippocampus is its capacity to sustain neurogenesis beyond birth^2,3^. To assess whether hippocampal organoids recapitulate this property, we examined NSC maintenance by SOX2 immunostaining from day 30 to day 90 **(Figure 2A)**, revealing organized NSC layers resembling those in the dentate neuroepithelium of the human fetal hippocampus at GW16 **(Figure 2B)**. Organoids displayed organized SOX2^+^ NSC layers, with layer thickness peaking at day 50 **(Figure 2C)**. The DCX^+^ immature neuron layer was also sustained throughout culture, peaking in thickness at day 70 **(Figure 2C-D)**, indicating ongoing neurogenesis.

**Figure 2.**
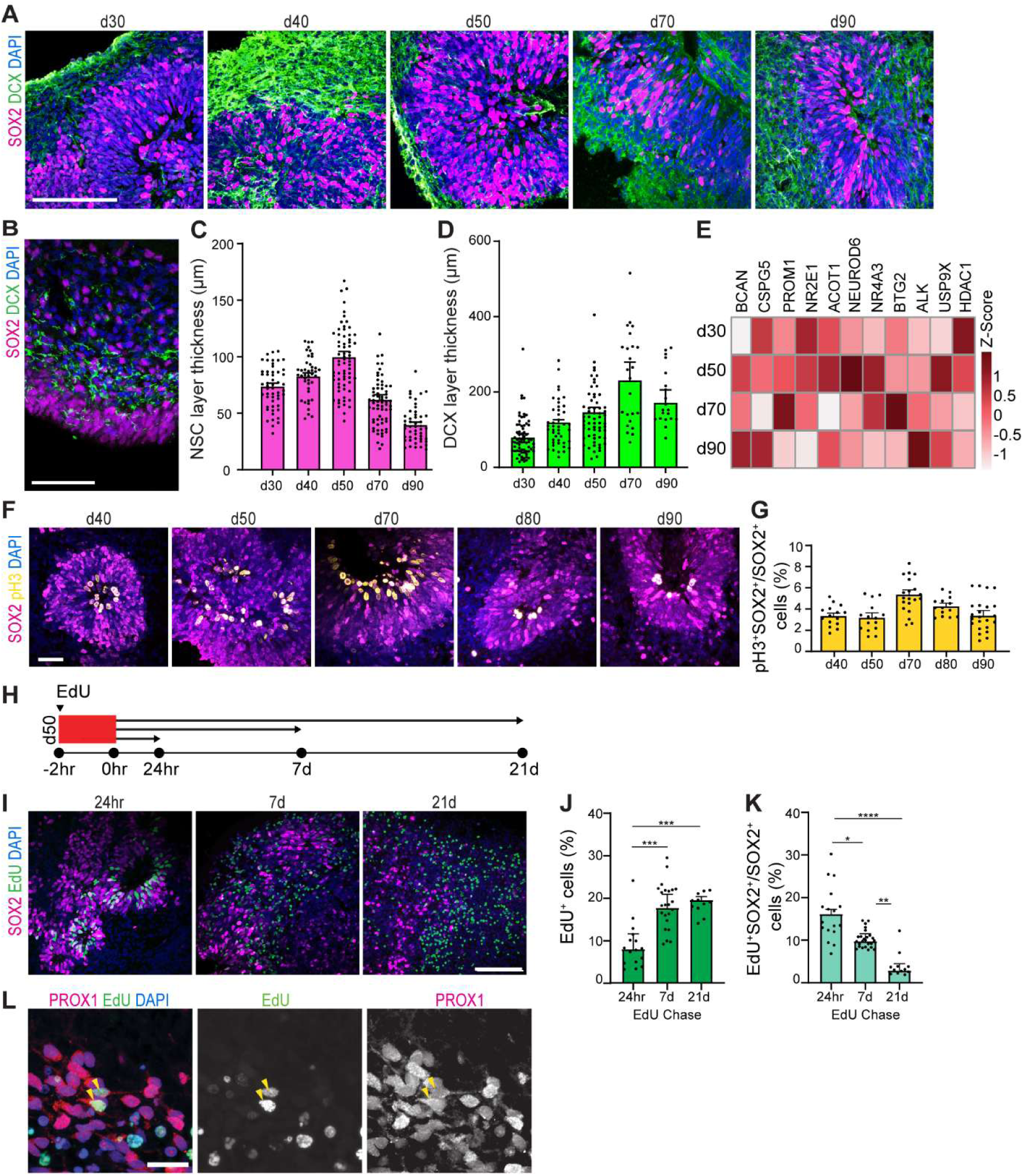
Maintenance of NSC population and proliferative activity over time in hippocampal organoids. **(A and B)** Representative immunofluorescence images showing SOX2^+^ NSCs (magenta) and DCX^+^ immature neurons (green) in hippocampal organoids d20-d90) (**A**) and GW16 fetal hippocampus (**B**). Scale bars = 100 μm. **(C and D)** Quantification of SOX2^+^ NSC layer thickness (d20-d90) (**C**) and DCX^+^ layer thickness (**D**) (d30-d90) (mean + SEM, n = 37-68 measurements from ≥3 organoids per timepoint). **(E)** Heatmap depicting the expression of genes associated with NSC maintenance and function, derived from RNA-sequencing analysis, across different stages of hippocampal organoid development. **(F)** Representative Immunofluorescence images of mitotic pH3^+^ (yellow) cells within SOX2^+^NSC layers (magenta) at d40-d90, localized at the ventricular side of the NSC layer. Scale bar = 50 μm. **(G)** Quantification of the proportion of pH3^+^SOX2^+^ cells shows ∼3-5% mitotically active NSCs from d40-d90 (mean + SEM, n = 15-23 images from ≥3 organoids per timepoint). **(H)** Schematic diagram of EdU pulse-chase paradigm: 2hr EdU pulse at d50 followed by 24hr, 7d, or 21d chase. **(I)** Representative immunofluorescence images displaying EdU^+^ (green) cells and SOX2^+^ (magenta) NSCs after a 24hr (left), 7d (middle) or 21d (right) chase period. Scale bar = 100 μm. **(J)** EdU^+^ cell numbers increase significantly from 24hr to 7d and 21d (Kruskal-Wallis, 24hr vs 7d: ***p=0.0001, 24hr vs 21d: ***p=0.0007; median ± IQR; n = 11-23 images from ≥3 organoids per timepoint). **(K)** Proportion of EdU^+^SOX2^+^ NSCs decreases over time, indicating NSC differentiation; Kruskal-Wallis, *p=0.02, **p=0.002, ****p<0.0001 median ± IQR, n = 12-25 images from ≥3 organoids per timepoint). **(L)** Representative immunofluorescence image of EdU^+^ PROX1^+^ dentate granule cells in hippocampal organoids (arrows). PROX1^+^ dentate granule cells shown in magenta, EdU^+^ cells shown in green. Scale bar = 20 μm.

Transcriptomic profiling revealed sustained expression of multiple genes associated with NSC maintenance, survival, and differentiation, including *PROM1*^36^, *BCAN*^37^, *CSPG5*^38^, *NR2E1*^39^, *ACOT1*^40,41^, and *NEUROD6*^42^. Additional markers such as *NR4A3*^43^, *BTG2*^44^, *ALK*, and *HDAC1*^45^ further supported active neurogenic processes, while continuous expression of USP9X, critical for hippocampal development and NSC self-renewal, highlighted the preservation of functional NSC populations **(Figure 2E)**. Together, these findings demonstrate that hippocampal organoids maintain organized NSC niches with ongoing neurogenic capacity throughout extended culture.

We next assessed NSC proliferative capacity by co-staining for SOX2 and pH3. Mitotically active NSCs (3-5%) were consistently observed across days 40-90 **(Figure 2F-G)**. To further probe neurogenic potential, EdU pulse-chase labeling was performed at day 50 **(Figure 2H)**. The proportion of EdU^+^ cells increased significantly from 8.88% after 24 hours to 18.32% at 7 days, reflecting active proliferation, before stabilizing at 21 days **(Figure 2I** and **J)**. Within SOX2^+^ NSCs, EdU labeling decreased over time, consistent with ongoing division and differentiation **(Figure 2I** and **K)**. Importantly, PROX1^+^EdU^+^ neurons were identified on day 21, confirming the generation of dentate granule cells from dividing NSCs **(Figure 2L)**. Together, these findings demonstrate that hippocampal organoids maintain proliferative NSCs and support sustained neurogenesis over extended culture periods.

### Progressive Neuronal Maturation and Synapse Formation in Hippocampal Organoids

Beyond sustaining self-renewing NSCs, it is critical that hippocampal organoids generate mature, functional neurons. To evaluate neuronal maturation, we performed immunohistochemistry for MAP2, a neuronal microtubule-associated protein. MAP2^+^ neurons progressively increased from day 30 to 90, forming complex networks with widespread expression across organoids **(Figure 3A)**. Additional markers of maturation were detected, including synaptophysin, a presynaptic vesicle glycoprotein, and NeuN, a nuclear antigen of mature neurons **(Figure 3B-E)**, indicating the establishment of synaptic potential. We also identified GAD67^+^ inhibitory neurons **(Figure 3F)** and VGAT, a GABAergic synaptic marker **(Figure 3G-H)**. Ultrastructural validation by transmission electron microscopy confirmed synapse formation, revealing pre- and post-synaptic compartments and vesicle accumulation **(Figure 3I)**. Together, these findings demonstrate that organoids generate diverse, mature neuronal populations with synaptic features.

**Figure 3.**
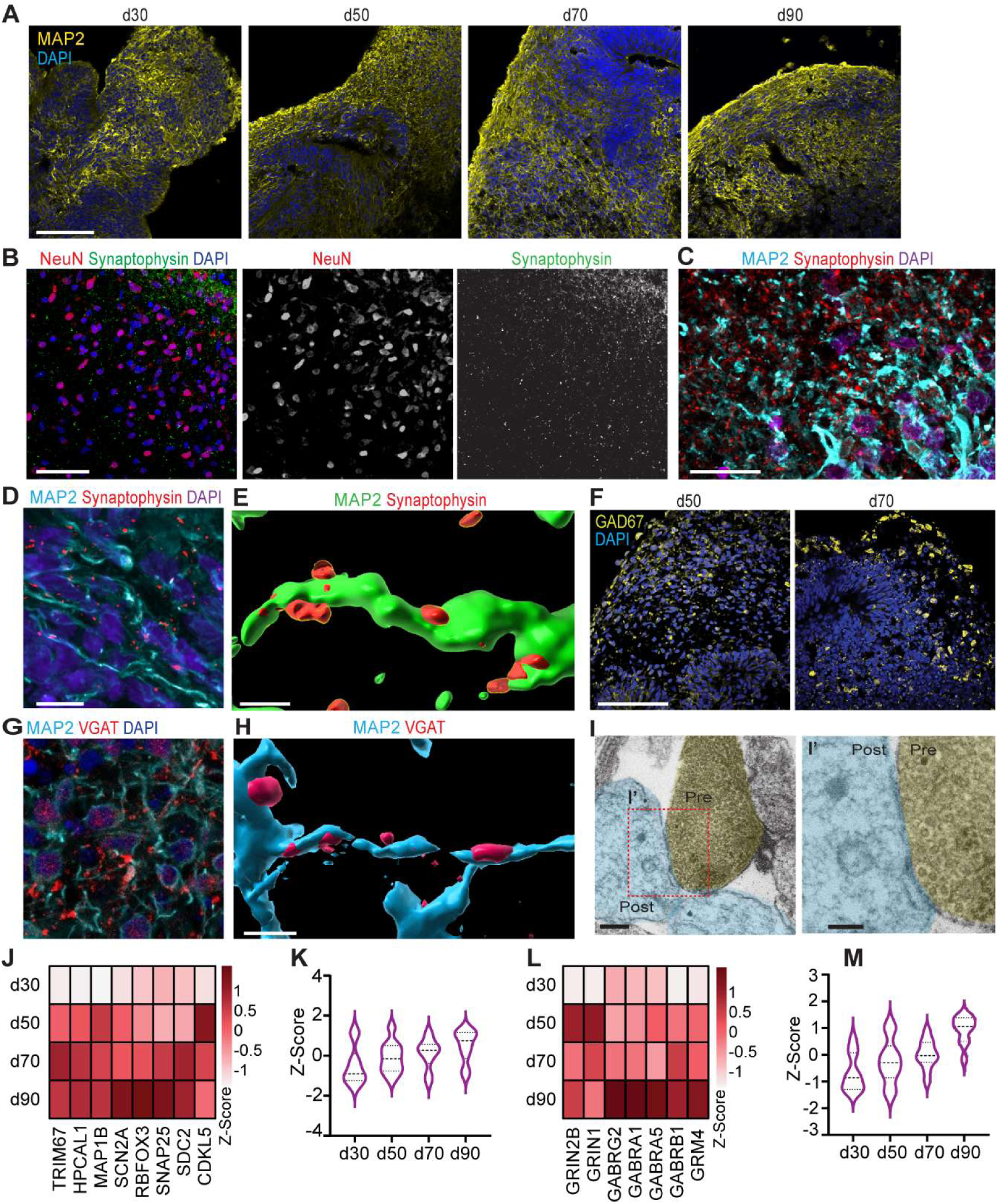
Neuronal maturation and synaptogenesis in hippocampal organoids. **(A)** Representative immunofluorescence images showing progressive increase of MAP2^+^ neurons (yellow) from day (d) 30 to d90. Scale bar = 100 μm. **(B)** Representative immunofluorescence images of mature neurons (NeuN^+^, red) and synapses (SYN, green) in a d90 organoid. Scale bar = 50 μm. **(C and D)** Representative maximum intensity projection images of neurons (MAP2^+^, blue) and SYN^+^ synapses (red) in GW16 fetal hippocampus (**C**) and d90 hippocampal organoid (**D**). Scale bar = 20 μm (**C**), 10 μm (**D**). (**E**) Representative 3D reconstruction image of MAP2^+^ neurons (green) and SYN^+^ synapses (red) from (**D**) using Imaris. Scale bar = 10 μm. (**F**) Representative immunofluorescence images of GAD67^+^ inhibitory neurons (yellow) in d50 (left) and d70 (right) hippocampal organoids. Scale bar = 100 μm. (**G**) Representative immunofluorescence images neurons (MAP2^+^, red) and inhibitory synapses (VGAT^+,^ blue) in d80 hippocampal organoid. Scale bar = 20 μm. (**H**) Representative 3D reconstruction image of MAP2^+^ neurons (blue) and VGAT^+^GABAergic synapses (red) from (**G**) using Imaris. Scale bar = 10 μm. (**I**) Transmission electron microscopy image of a synapse in d90 organoid; pseudo colored pre-(yellow) and post-synaptic (blue) areas. Scale bars: 500 nm (**I**), 100 nm (**I’**). **(J and K)** Heatmap (**J**) and violin plot (**K**) depicting the expression of genes associated with neuronal maturation and synapse development, derived from RNA-sequencing analysis, across different stages of hippocampal organoid development (d30-d90). **(L and M)** Heatmap (**L**) and violin plot (**M**) depicting the expression of genes associated with excitatory and inhibitory neurotransmission, derived from RNA-sequencing analysis, across different stages of hippocampal organoid development (d30-d90).

Transcriptomic profiling further supported progressive neuronal maturation. Across days 30-90, RNA sequencing revealed increased expression of genes involved in neuronal differentiation, synapse formation, and neurotransmission **(Figure 3J-M)**. *RBFOX3* (NeuN) and *SNAP25*, critical for synaptic function, were progressively upregulated, consistent with advancing neuronal maturation. *SCN2A*, encoding the sodium channel Nav1.2 essential for action potential initiation, also increased, reflecting electrophysiological development. Genes linked to neuronal structure and synaptogenesis, including *MAP1B* and *SDC2*, were similarly upregulated, aligning with dendritic spine formation and maturing neuronal networks **(Figure 3J)**. Collectively, neuronal maturation gene expression shifted markedly, with average Z-scores rising from −0.51 at day 30 to 0.50 by day 90 **(Figure 3K)**. In contrast, *CDKL5* expression declined **(Figure 3J)**, consistent with its developmental downregulation in postnatal brain, further indicating maturation.

Expression of NMDA and GABA receptor subunits (*GRIN2B*, *GRIN1*, *GABRG2*, *GABRA1*, *GABRA5*, *GABRB1*) increased over time, supporting the establishment of balanced excitatory and inhibitory synaptic transmission **(Figure 3L)**. The average Z-score for synaptic genes rose from −0.6 at day 30 to 0.9 at day 90 **(Figure 3M)**, reinforcing the progressive development of functional neuronal networks.

Together, these results demonstrate that hippocampal organoids not only sustain NSCs but also undergo physiologically relevant neuronal maturation, establishing synaptic architecture and functional potential over extended culture.

### Functional Maturation and Electrophysiological Activity in Hippocampal Organoids

Spontaneous neuronal activity in the developing brain is initially driven by calcium fluctuations, which support maturation and circuit formation. To assess this in hippocampal organoids, spontaneous calcium transients were measured using Fluo-4 AM across developmental stages **(Figure 4A)**. At day 30, sparse activity was detected, but by day 50 both the number of active neurons and the amplitude of transients increased significantly **(Figure 4B-D)**. Neurons at both stages displayed synchronous firing, indicating early network formation **(Figure 4E)**. By day 70, activity became spatially organized, with distinct groups of neurons firing synchronously, consistent with emerging microcircuitry. The number of active neurons, amplitude of calcium transients, and cumulative activity continued to rise, supporting progressive maturation of neuronal excitability and network coordination **(Figure 4B-E)**.

**Figure 4.**
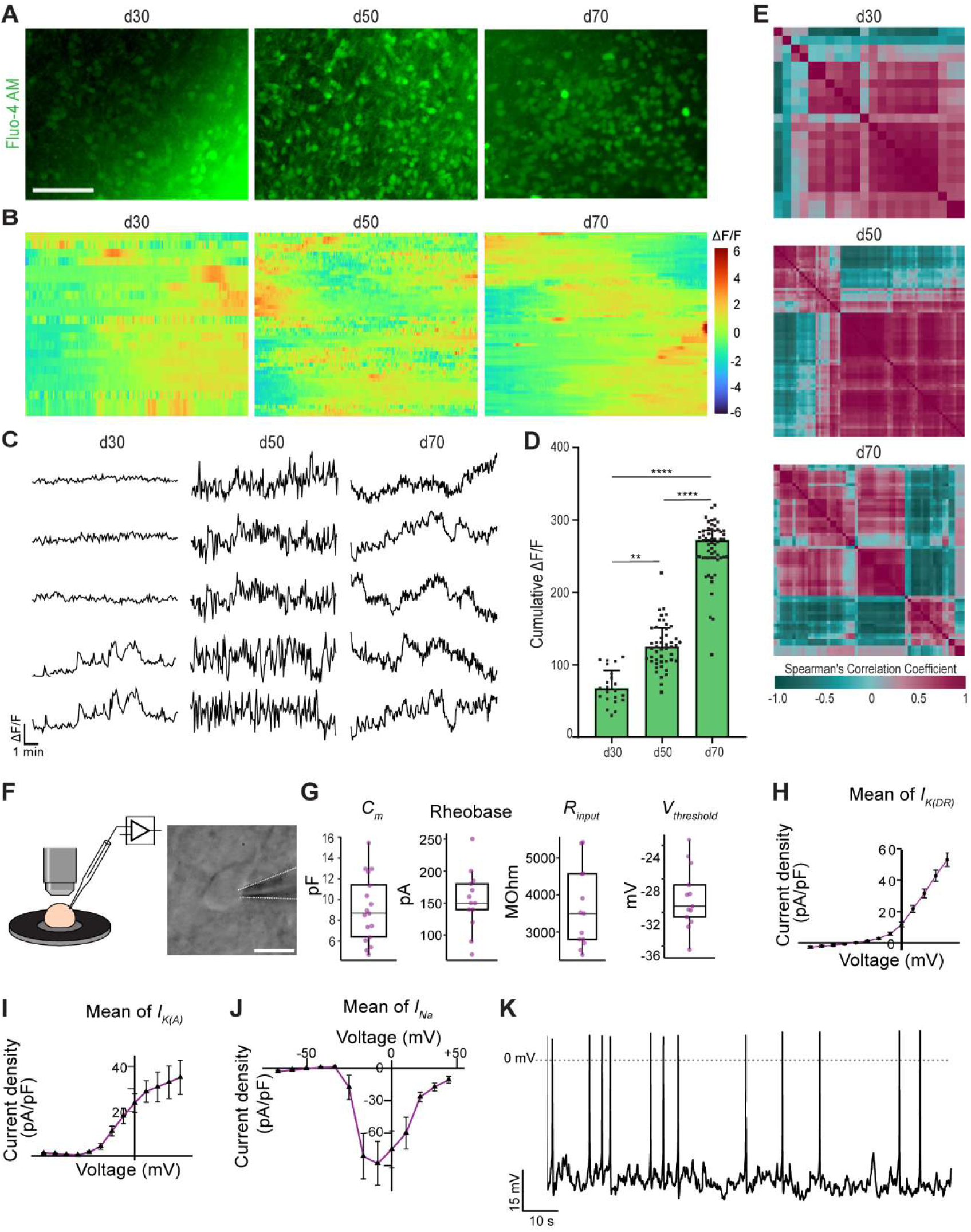
Functional maturation and electrophysiological properties of hippocampal organoid neurons. **(A)** Representative Immunofluorescence images of Fluo-4 AM (green) in hippocampal organoids at day (d) 30, d50, and d70. Scale bar = 200 μm. **(B)** Heat maps of ΔF/F calcium signals in individual neurons over time (d30-d70), showing increased frequency and synchrony with maturation. **(C)** Representative calcium transient traces at d30, d50, and d70. **(D)** Quantification of cumulative ΔF/F values indicating increased calcium activity over time (Kruskal-Wallis, **p=0.005, ****p<0.0001; median ± IQR; n = 22-58 neurons from 9 organoids). **(E)** Correlation matrices showing transition from sparse to organized synchronous firing from d30 to d70. **(F)** Schematic (left) and representative image (right) of whole-cell patch clamp setup with recording pipette visible. **(G)** Intrinsic membrane properties (capacitance, rheobase, input resistance, action potential threshold) measured in neurons from d70-d160 hippocampal organoids (median ± IQR; n = 13-18 neurons from 9 organoids). **(H-J)** Voltage-dependent I_K(DR)_, I_K(A)_, and I_Na_ current-voltage relationships showing typical neuronal ion channel activity (mean ± SD; n = 9-14 neurons from 9 organoids). **(K)** Whole-cell current-clamp trace showing spontaneous action potential firing in a d90 neuron.

To further assess functional properties, whole-cell patch-clamp recordings were performed on hippocampal organoids between days 70-160 **(Figure 4F)**. Neurons at the organoid periphery displayed intrinsic electrical properties consistent with neuronal maturation, including a mean membrane capacitance of 9.1 pF, rheobase of 154.2 pA, input resistance of 3.6 GΩ, and an action potential threshold of −29.4 mV **(Figure 4G)**. These values align with previously reported parameters for neurons in brain organoids, confirming physiological relevance^46,47^.

Voltage-clamp experiments revealed the presence of multiple ion channel currents. Two outward potassium currents were identified: the delayed rectifier potassium current, *I_K(DR)_*, and transient A-type current, *I_K(A)_*, isolated using subtraction protocols **(Figure S4A-D)**. *I_K(DR)_* activated near −30 mV and reached ∼50 pA/pF at +50 mV **(Figure 4H)**, while *I_K(A)_* showed transient activation with peak current densities of ∼35 pA/pF at +50 mV **(Figure 4I)**. The detection of both currents indicates neuronal heterogeneity and supports the presence of maturing subtypes with distinct repolarization dynamics. Inward sodium currents, *I_Na_*, recorded with voltage steps **(Figure S4E)**, exhibited transient activation beginning near −30 mV, peaking at −10 mV with ∼-90 pA/pF, and rapidly inactivating at more depolarized potentials **(Figure 4J)**. This profile is consistent with fast-activating sodium channels required for action potential initiation.

Importantly, neurons within the organoids were capable of firing action potentials. Representative traces from day 90 organoids revealed spontaneous trains of spikes with consistent waveforms, confirming the coordinated activity of sodium and potassium channels **(Figure 4K)**.

Together, patch-clamp and ion current analyses demonstrate that hippocampal organoid neurons acquire functional electrophysiological properties, including active sodium and potassium conductance and the ability to fire spontaneous action potentials. These findings provide strong evidence of progressive neuronal maturation and validate the organoids as a model for functional human hippocampal circuitry.

### Astrogenesis and Long-Term Stability of Hippocampal Organoids

During hippocampal development, astrogenesis follows neurogenesis making the emergence of astrocytes a key indicator that organoids are recapitulating a physiologically relevant developmental trajectory^48^. Astrocyte generation within hippocampal organoids was assessed by immunohistochemistry and transcriptomics. GFAP^+^ and S100β^+^ astrocytes were detected, predominantly at the periphery, displaying characteristic astrocytic morphology **(Figure 5A** and **B)**. Between day 70 and day 110, the percentage of GFAP^+^ area increased, suggesting ongoing astrogenesis **(Figure 5C** and **D)**. Transcriptomic profiling further supported these findings, showing reduced expression of immature astrocytic markers (*FGFR3*, *BMPR1B*, *SLC1A3*) and increased expression of mature astrocytic markers (*GFAP*, *S100β*, *SLC4A4*, *PAQR6*, *SLC1A2*), with Z-scores shifting accordingly between day 30 and day 90 **(Figure 5E-G)**. The presence of mature astrocytes is consistent with human hippocampal development, where astrocytes contribute to synaptic organization, neurotransmitter homeostasis, and neuronal metabolic support, enhancing the physiological relevance of the model^49,50^.

**Figure 5.**
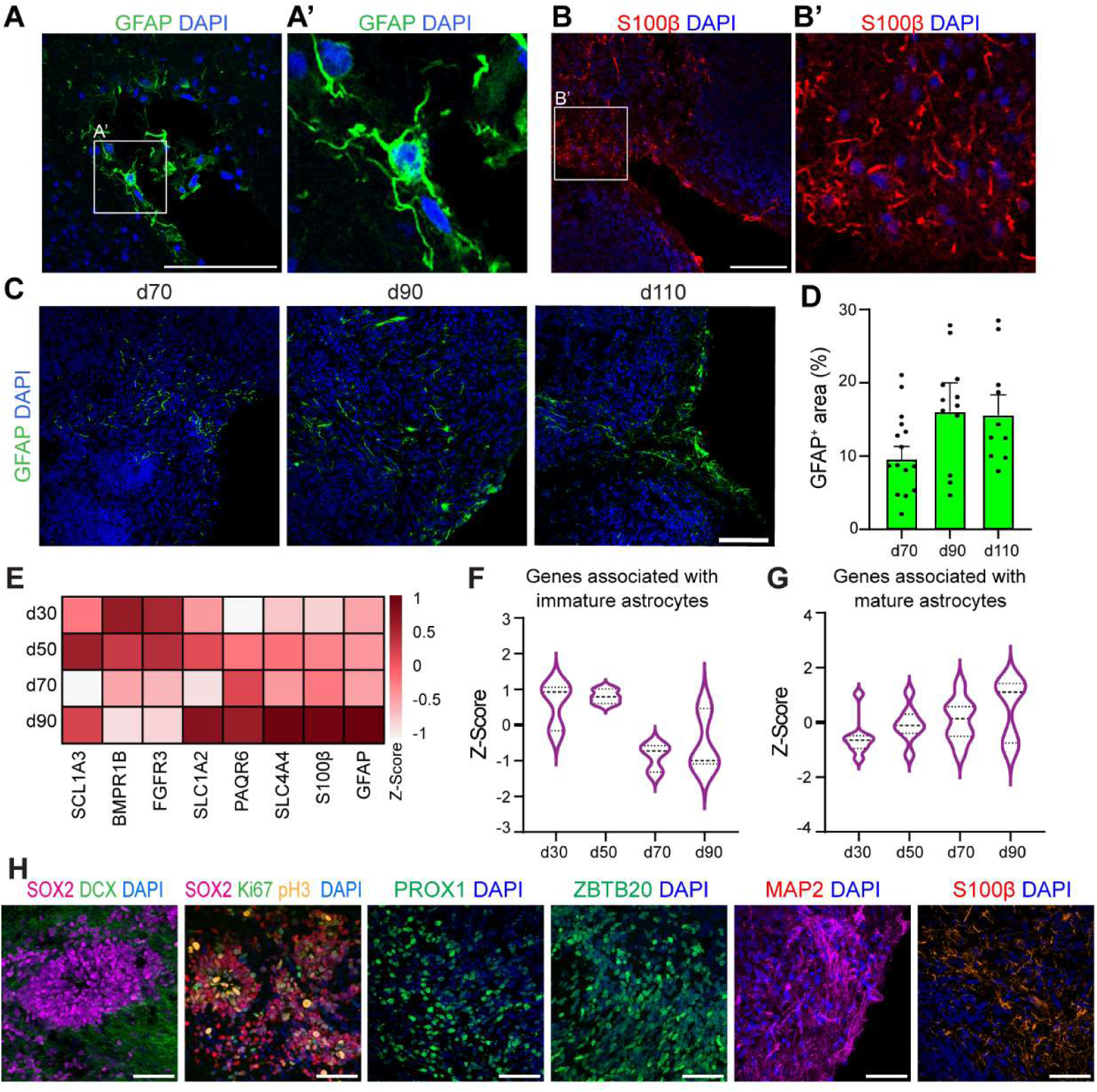
Astrogenesis and Long-Term Stability of Hippocampal Organoids. **(A and B)** Representative immunofluorescence images of astrocytes showing GFAP (green, **A**) and S100β (red, **B**) expression with characteristic astrocytic morphology (**A′, B**′) at the organoid periphery at day 90. Scale bar, 100 μm **(C and D)** Progressive increase in GFAP^+^ astrocytes from d70 to d110 (**C**), confirmed by quantification of GFAP^+^ area (**D**) (mean + SEM; n = 7-15 images from ≥3 organoids per timepoint). **(E-G)** Heatmap (**E**) and violin plots (**F, G**) depicting the expression of genes associated with astrocytes development (immature astrocyte (**F**) and mature astrocytes (**G**), derived from RNA-sequencing analysis, across different stages of hippocampal organoid development (d30-d90). **(H)** Representative immunofluorescence images d120 organoids shows radial growth trends with maintenance of NSC layers (SOX2, magenta), immature neurons (DCX, green), proliferating (Ki67, green) and mitotic (pH3, yellow) NSCs, dentate granule cells (PROX1, green), hippocampal neurons (ZBTB20, green), neurons (MAP2, red), and astrocytes (S100β, red). Scale bars = 50 μm.

In addition to astrocyte generation, hippocampal organoids demonstrated long-term stability in culture, with sustained growth and cell-type maintenance beyond 120 days **(Figure 5H)**. Organoids preserved SOX2^+^ NSC niches, maintained mitotic activity, and continued to generate DCX^+^ immature neurons, confirming ongoing neurogenesis at late stages.

Neuronal populations, including PROX1^+^ dentate granule cells, ZBTB20^+^ hippocampal neurons, and MAP2^+^ neurons, were also retained. In parallel, S100β^+^ astrocytes persisted, ensuring the coexistence of NSCs, neurons, and astrocytes over extended culture.

Together, these findings demonstrate that hippocampal organoids robustly recapitulate the sequential generation and long-term maintenance of key hippocampal cell types. Their ability to sustain organized NSC niches, neuronal populations, and astrocytes for months in culture provides a powerful platform for studying extended developmental timelines, neurogenesis, and chronic disease processes in vitro.

### Glucocorticoid Signaling Drives NSC Quiescence in Hippocampal Organoids

Glucocorticoids are key regulators of stress responses and fetal brain development^51^, but excessive exposure, such as through maternal stress or prenatal corticosteroid therapy, has been linked to hippocampal impairment in animal and human studies^14–24^. To investigate the effect of excessive glucocorticoid signaling on human hippocampal development with a particular focus on NSCs, organoids were treated with dexamethasone, a synthetic glucocorticoid, at days 44, 46, and 48, followed by analysis at day 50, a stage marked by peak NSC layer thickness **(Figure 6A)**.

**Figure 6.**
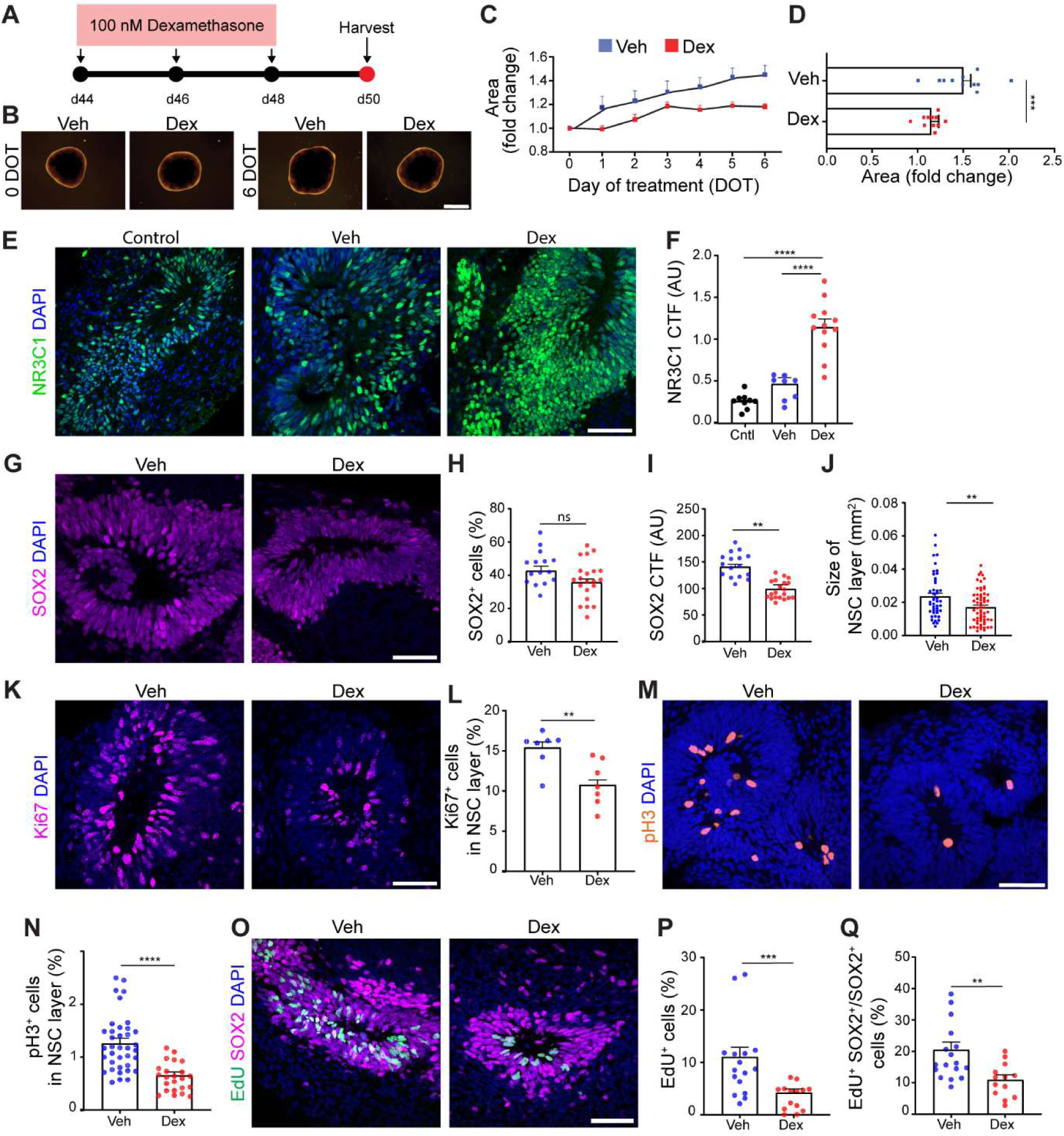
Dexamethasone treatment reduces hippocampal organoid growth and neural stem cell proliferation. **(A)** Experimental timeline: hippocampal organoids were treated with 100 nM dexamethasone (Dex) or vehicle (Veh) at d44, d46, and d48, followed by analysis at d50 **(B and C)** Representative brightfield images (**B**) and growth patterns (**C**) in Veh and Dex show reduced expansion of dexamethasone-treated organoids over 6-day treatment paradigm. Scale bar = 1 mm. n = 11 organoids per group. **(D)** Quantification of overall change in organoid area confirms significant growth reduction after dexamethasone treatment (Unpaired t-test, ***p=0.0007; mean + SEM; n = 11 organoids at d44 and d50). **(E and F)** Representative immunofluorescence images of NR3C1 expression (**E**, green) and quantification (**F**) in control, Veh and Dex. Scale bar = 50 µm (Unpaired t-test, ****p<0.0001; mean + SEM; n = 9-12 images from 3 organoids at d50). **(G-J)** Representative immunofluorescence image of SOX2^+^ NSC in Veh and Dex (**G**, magenta) and quantifications of number of SOX2^+^ cell (**H**), SOX2 intensity (**I**) and NSC layer size (**J**). Scale bar = 50 µm (Unpaired t-test, **p=0.005, ***p=0.0002; mean + SEM; n = 16-61 images from 3 organoids at d50). **(K-N)** Representative immunofluorescence image of proliferating (**K**, Ki67^+^, magenta) and mitotic (**M**, pH3^+^, orange) NSCs in Veh and Dex and their quantifications (**L, N**). Scale bar = 50 µm (Mann-Whitney test, **p=0.007, ****p<0.0001; mean + SEM; n = 7-37 images from 3 organoids at d50). **(O-Q)** Representative immunofluorescence images of EdU^+^ cells in NSC layer in Veh and Dex (**O**) and quantifications of EdU^+^ proliferating cells (**P**) and EdU^+^SOX2^+^ NSCs (**Q**). Scale bar = 50 µm (Mann-Whitney test, **p=0.008, ***p=0.0009; mean + SEM; n = 13-17 images from 3 organoids at d50).

Brightfield imaging revealed no overt morphological abnormalities upon dexamethasone treatment, but quantitative analysis showed that dexamethasone-treated organoids exhibited significantly reduced growth compared with vehicle controls. By day 6 of treatment, vehicle-treated organoids increased 1.5-fold in size, whereas dexamethasone-treated organoids expanded only 1.15-fold (p = 0.0007) **(Figure 6B-D)**. Immunostaining demonstrated a robust upregulation and increased nuclear localization of the glucocorticoid receptor NR3C1 within NSC layers upon dexamethasone treatment, confirming effective NR3C1 pathway activation **(Figure 6E-F)**.

To investigate whether the observed reduction in organoid growth was associated with altered NSC dynamics, NSC layers were assessed following dexamethasone exposure. SOX2 staining showed that fluorescence intensity and NSC layer thickness were significantly reduced in dexamethasone-treated organoids while NSC numbers were unchanged **(Figure 6G-J)**, suggesting increased density of NSC layers potentially due to altered NSC state or transcriptional activity. Functional assays showed reduced proliferation, with fewer Ki67^+^ NSCs and nearly half the number of mitotic pH3^+^ NSCs **(Figure 6K-N)**. EdU pulse-chase experiments further confirmed these findings, revealing a marked reduction in proliferating NSCs, with EdU^+^ SOX2^+^ cells falling from 19.2% to 10.8% **(Figure 6O-Q)**.

To explore the molecular mechanisms underlying altered NSC dynamics, we performed bulk RNA-sequencing on dexamethasone-treated hippocampal organoids. Differentially expressed genes (DEGs) were enriched in pathways related to autophagy, DNA repair, and senescence **(Figure S5)**. Notable DEGs included *SYNGAP1*, a regulator of MAPK/ERK signaling that influences NSC proliferation and survival^52,53^, histone H4 variants *H4C12* and *H4C8*, linked to chromatin remodeling^54^, and *RAD50*, a DNA repair factor^55^ **(Figure S5A)**. These findings suggest dexamethasone activates stress-response pathways that may influence NSC state.

Reduced NSC proliferation may reflect a shift into quiescence, a defining feature of hippocampal NSCs that preserves their proliferative potential and sustains neurogenesis after birth^56,57^. Yet, the molecular mechanisms governing the transition from active proliferation to quiescence remain only partially understood. To investigate whether glucocorticoid receptor activity contributes to NSC quiescence, we examined previously published transcriptomic datasets from two studies by Berg et al. and Jimenez-Cyrus et al.^6,58^, which profiled gene expression during the transition of hippocampal NSCs from active to quiescent states. In both datasets, expression of the glucocorticoid receptor gene NR3C1 was significantly upregulated as NSCs entered quiescence **(Figure S6A and B)**. Given the observed increase in NR3C1 expression in quiescent NSCs, we hypothesized that dexamethasone promotes a quiescent-like state in hippocampal organoids. To test this, we examined gene modules identified in the study by Jimenz-Cyrus et al.^58^ that define the transition from active to quiescent NSCs. These include “off” genes downregulated during quiescence and “on” genes upregulated as cells enter this state. Dexamethasone-treated organoids showed a trend toward reduced expression of “off” genes, supporting a loss of active NSC identity, while “on” genes exhibited a trend toward increased expression **(Figure 7A-C)**. Notably, *BHLHE41*, a regulator of hippocampal NSC quiescence, and additional markers such as *ETV5*, *EPAS1*, *HOPX*, and *HES1*, all increased in expression, reinforcing the hypothesis that glucocorticoids shift NSCs toward quiescence.

**Figure 7.**
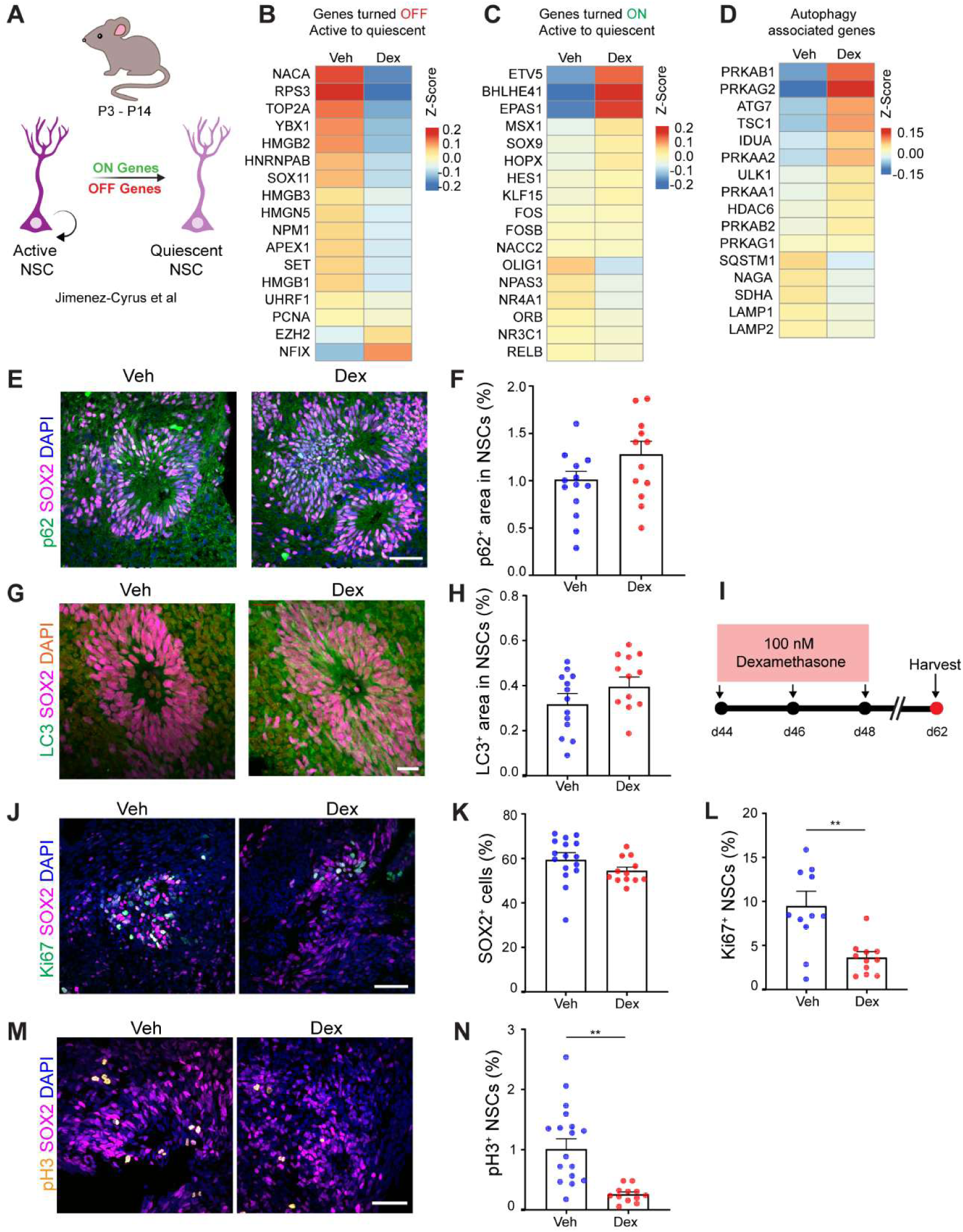
Dexamethasone treatment drives transcriptional and cellular signatures of NSC quiescence in hippocampal organoids. **(A)** Schematic summary of NSC transition from an active to a quiescent state, adapted from Jimenez-Cyrus et al. (2024). The study identified two gene modules: genes downregulated (“off”) and genes upregulated (“on”) during the transition of dentate gyrus NSCs between postnatal days 3 and 14 (P3-P14) in mice. **(B and C)** Heatmaps showing expression of “off” (**B**) and “on” (**C**) gene modules associated with the transition of NSCs from an active to a quiescent state. In hippocampal organoids, dexamethasone treatment (Dex) reduced expression of “off” genes and increased expression of “on” genes compared to vehicle (Veh). Gene sets were obtained from Jimenez-Cyrus et al., 2024 (n = 3 organoids per group). **(D)** Heatmap showing expression of autophagy-related genes associated with NSC quiescence (Calatayud-Baselga et al., 2023), which are upregulated in hippocampal organoids following dexamethasone treatment (n = 3 organoids per group). (**E and F**) Representative immunofluorescence images of p62 immunoreactivity (green) in SOX2^+^ NSC layer (magenta) in Veh and Dex (**E**) and quantifications (**F**). Scale bar = 50 µm (Unpaired t-test, p = 0.08; mean + SEM; n = 12-13 images from 3 organoids at d50). **(G and H)** Representative immunofluorescence images of LC3 immunoreactivity (green) in SOX2^+^ NSC layer (magenta) in Veh and Dex and their quantifications (**H**). Scale bar = 50 µm (Unpaired t-test, p=0.06; mean + SEM; n = 12-13 images from 3 organoids). **(I)** Experimental paradigm to examine NSC quiescence maintenance: 100 nM dexamethasone given at d44, d46, d48; analysis at d62 (14 days post-treatment). **(J-L)** Representative immunofluorescence images of proliferating Ki67^+^SOX2^+^ NSCs) in Veh and Dex (**J**) and their quantifications (**K** and **L**). Scale bar = 50 µm Unpaired t-test, **p=0.0013; mean + SEM; n = 11-17 images from 4 organoids at d62). **(M-N)** Representative immunofluorescence images of mitotic pH3^+^SOX2^+^ NSCs in Veh and Dex (**M**) and their quantifications (**N)**. Scale bar = 50 µm (Unpaired t-test, ****p<0.0001, mean + SEM; n = 12-17 images from 4 organoids at d62).

Prior studies have shown that autophagy is required for the developmental transition from proliferative to quiescent NSCs^58,59^. In dexamethasone-treated organoids, RNA-sequencing revealed a trend of increased expression of key autophagy genes, including AMPK subunits (*PRKAB1*, *PRKAG2*, *PRKAA1*, *PRKAB2*), *ULK1*, *TSC1*, and *ATG7*, which regulates autophagosome formation and has been shown to be essential for NSC quiescence in vivo^59–62^. Some late-stage autophagy and lysosomal genes such as *LAMP1*, *LAMP2*, *SQSTM1* were modestly reduced, possibly reflecting pathway timing or the averaging effects of bulk RNA-sequencing **(Figure 7D)**.

At the cellular level, immunostaining supported these transcriptomic findings. Both p62 and LC3, markers of autophagic flux and autophagosome formation respectively, showed a consistent trend toward increased abundance within SOX2^+^ NSC layers after dexamethasone exposure **(Figure 7E-H)**.

Finally, we assessed whether dexamethasone-induced quiescence persisted beyond the treatment window. Organoids were treated with dexamethasone on days 44-48 and analyzed two weeks post the last date of treatment **(Figure 7I)**. Dexamethasone treated organoids maintained comparable proportions of SOX2^+^ NSCs relative to controls **(Figure 7J** and **K)**, indicating preservation of the stem cell pool. However, Ki67^+^ and pH3^+^ NSCs were markedly reduced **(Figure 7L-N)**, demonstrating a lasting suppression of NSC proliferation. This indicates that glucocorticoid exposure drives NSCs into a stable quiescent-like state without depleting their numbers, paralleling in vivo dynamics where NSC pools are maintained despite reduced proliferative activity^63^. Importantly, the reduced proliferation of NSCs was not attributable to senescence. β-galactosidase staining, p21 expression, and γH2AX foci analysis showed no increase in senescent cells or DNA damage **(Figure S6C-K)**. These findings indicate that dexamethasone suppresses NSC proliferation without engaging canonical senescence pathways, instead promoting a quiescent-like state.

Together, these data reveal that dexamethasone enforces a premature transition of hippocampal NSCs into quiescence-like state and activation of autophagy-associated programs, suggesting a mechanism by which prenatal glucocorticoid exposure may impair hippocampal growth while preserving latent neurogenic capacity.

## Discussion

In this study, we establish an optimized human iPSC-derived hippocampal organoid system that recapitulates key aspects of human hippocampal development. The organoids preserve well organized NSC layers that sustain active neurogenesis, generate hippocampus-specific neuronal cell types, and produce functional neurons, all confirmed at cellular and transcriptomic levels. They also display transcriptomic signatures aligning with late fetal and early postnatal hippocampus, highlighting their relevance for modeling the prolonged developmental plasticity of this region. This framework provides a robust platform to investigate the pathways that regulate human hippocampal development and maturation. We further demonstrate the usefulness of this model by examining how glucocorticoid exposure alters neurogenesis, showing that stress hormone signalling disrupts typical neurogenic activity and promotes a shift of NSCs toward a quiescent-like state.

During hippocampal development, neurogenesis is confined to discrete germinal zones, including the dentate and hippocampal neuroepithelia, from which hippocampal cells are generated ^6,64^. A central advancement of this work is the refinement of differentiation conditions to yield radially organized organoids with well-defined, laminated NSC compartments that persist for months. Prior 2D systems and unguided cerebral organoids generated hippocampal-like neurons but did not report durable, structured NSC niches suitable for modeling hippocampal biology^25–27^. Likewise, hippocampal spheroids and early organoid protocols showed PROX1⁺ and ZBTB20⁺ neurons without demonstrating long-term NSC maintenance or organized progenitor zones, limiting evidence for sustained neurogenesis^28–30^. In contrast, our organoids establish clear NSC layers with radial polarity that continuously support ongoing neuronal production and gliogenesis. To improve reproducibility across iPSC lines with different proliferation rates and handling sensitivities, we optimized on EB size rather than cell number, identifying a size window that reliably produced radial polarity when combined with tuned WNT and BMP timing. This size-guided, timing-controlled protocol performed robustly across multiple iPSC lines, supporting its generalizability as a reproducible platform for modeling human hippocampal development.

The organoids also capture essential aspects of dentate gyrus NSC dynamics. In vivo, NSCs expand during embryonic development to generate PROX1⁺ granule neurons and later transition into a quiescent radial glia-like state that retains the capacity for reactivation^6,65^.

The organoids mirror this trajectory by maintaining organized NSC compartments that can switch between proliferative and dormant states in response to environmental cues. EdU pulse-chase labeling confirmed production of PROX1⁺ granule neurons from dividing NSCs, and comparative transcriptomics showed that organoids track the fetal hippocampal trajectory and most closely resemble late gestation to early postnatal hippocampus. This reflects the extended developmental plasticity of the hippocampus compared with other brain regions and highlights the utility of this model for dissecting NSC regulation in a human-specific context, where much of our current knowledge is otherwise inferred from rodent studies.

Using this platform, we found that dexamethasone reduces proliferation and shifts hippocampal NSCs toward a quiescent-like state while preserving overall NSC numbers. RNA-sequencing and immunostaining revealed activation of autophagy pathways within the organoids, including upregulation of AMPK-ULK1-ATG7 modules and increased LC3 and p62 expression. Glucocorticoid-induced autophagy has been reported in hippocampal NSCs, where corticosterone exposure promotes autophagy-associated cell death in adult mouse brain^66–68^, although this phenomenon has not been described during developmental stages in the human brain. Notably, autophagy has been implicated as a mechanism driving the developmental transition of hippocampal NSCs into quiescence, raising the possibility that glucocorticoids may co-opt this physiological program to enforce precocious dormancy.

While such a response may serve to protect the stem cell pool, if engaged prematurely, for example, under maternal stress or antenatal corticosteroid therapy, it could disturb NSC behavior and impair circuit maturation during critical developmental windows.

These findings carry broader implications within the developmental origins of health and disease framework, which emphasizes how maternal conditions and the intrauterine environment shape long-term health outcomes. Our data suggests that excessive glucocorticoid exposure can alter hippocampal developmental trajectories by prematurely shifting NSCs into quiescence, providing a potential cellular mechanism linking maternal stress to increased neurodevelopmental vulnerability in offspring. Beyond maternal stress, the model enables interrogation of how other environmental factors, such as infection, nutrition, or hypoxia, interfere with hippocampal development. The ability to sustain environmentally responsive NSCs in a human context also creates translational opportunities for screening interventions, testing candidate therapeutics, and studying disease mechanisms using patient-derived iPSCs.

Looking ahead, this platform can be extended to more faithfully model human hippocampal biology. Incorporating microglia and vascular components would enable investigation of neuroimmune and neurovascular interactions that shape development and plasticity. Fusion into assembloids, particularly with entorhinal cortex-like organoids, could reconstruct long-range inputs such as the perforant pathway and permit analysis of activity-dependent processes, including long-term potentiation. In parallel, single-cell and spatial transcriptomics will be essential for mapping lineage trajectories and revealing rare or transient states.

Together, these advances position human hippocampal organoids to address questions in human neurodevelopment that remain inaccessible in animal models and to bridge developmental biology, disease modeling, and translational neuroscience.

### Limitations of the study

The optimized protocol yields hippocampal organoids with sustained NSCs and ongoing neurogenesis, but several constraints remain. First, the organoids do not reproduce the characteristic V-shaped dentate gyrus or the displacement of NSCs away from the ventricular surface, so the in vivo germinal niche architecture is only partially modeled. Second, key niche components are absent, including vasculature and microglia, which influence neurogenesis, quiescence, and maturation. Third, the glucocorticoid paradigm relies on in vitro dexamethasone exposure and does not capture the timing, placental mediation, and systemic endocrine dynamics of prenatal stress or antenatal corticosteroid therapy. Finally, validation across diverse genetic backgrounds was constrained by the number of iPSC lines and biological replicates. Future work that incorporates vascular and immune cells, spatially patterned morphogens to promote dentate morphogenesis, assembloids to restore long-range inputs, and multi-line benchmarking will be important to address these gaps.

## Acknowledgments

We thank the members of the Kang and Berg laboratories for their valuable comments and suggestions. We are also grateful to the Fowler Lab team for their support with fetal sample acquisition. This work was supported by grants from the Academy of Medical Sciences (SBF007\100169 to E.K.), Medical Research Council (MR/Z506138/1 to E.K.), Alzheimer’s Society (636 to E.K.), Alzheimer’s Research UK (ARUK-PPG2024-029 to E.K.), The Royal Society (RG\R2\232559 to D.A.B.), and Biotechnology and Biological Sciences Research Council (BB/W008068/1 to D.A.B.). The SAFeR study was funded by the UK Medical Research Council (MR/L010011/1 and MR/P011535/1) and the EU’s Horizon 2020 research and innovation programme under the Marie Skłodowska-Curie project PROTECTED (grant agreement number 722634), as well as the FREIA and INITIALISE projects (grant agreement numbers 825100 and 101094099, respectively) awarded to P.A.F

## Author Contributions

E.K., D.A.B. and O.S. conceived the project and wrote the paper. O.S. led and contributed to all aspects of the study. M.Z.K.A. performed RNA-sequencing data analysis. J.A.G. carried out electrophysiology recordings and analysis. G.C. supported organoid culture. D.R. contributed to immunofluorescence staining of human fetal tissue. R.G. supported organoid culture and contributed to sectioning of organoids. P.A.F., S.S.M.V, and P.L. and L.L.L. contributed to obtaining fetal brain samples. All authors commented on and approved publication of manuscript.

## KEY RESOURCES TABLE

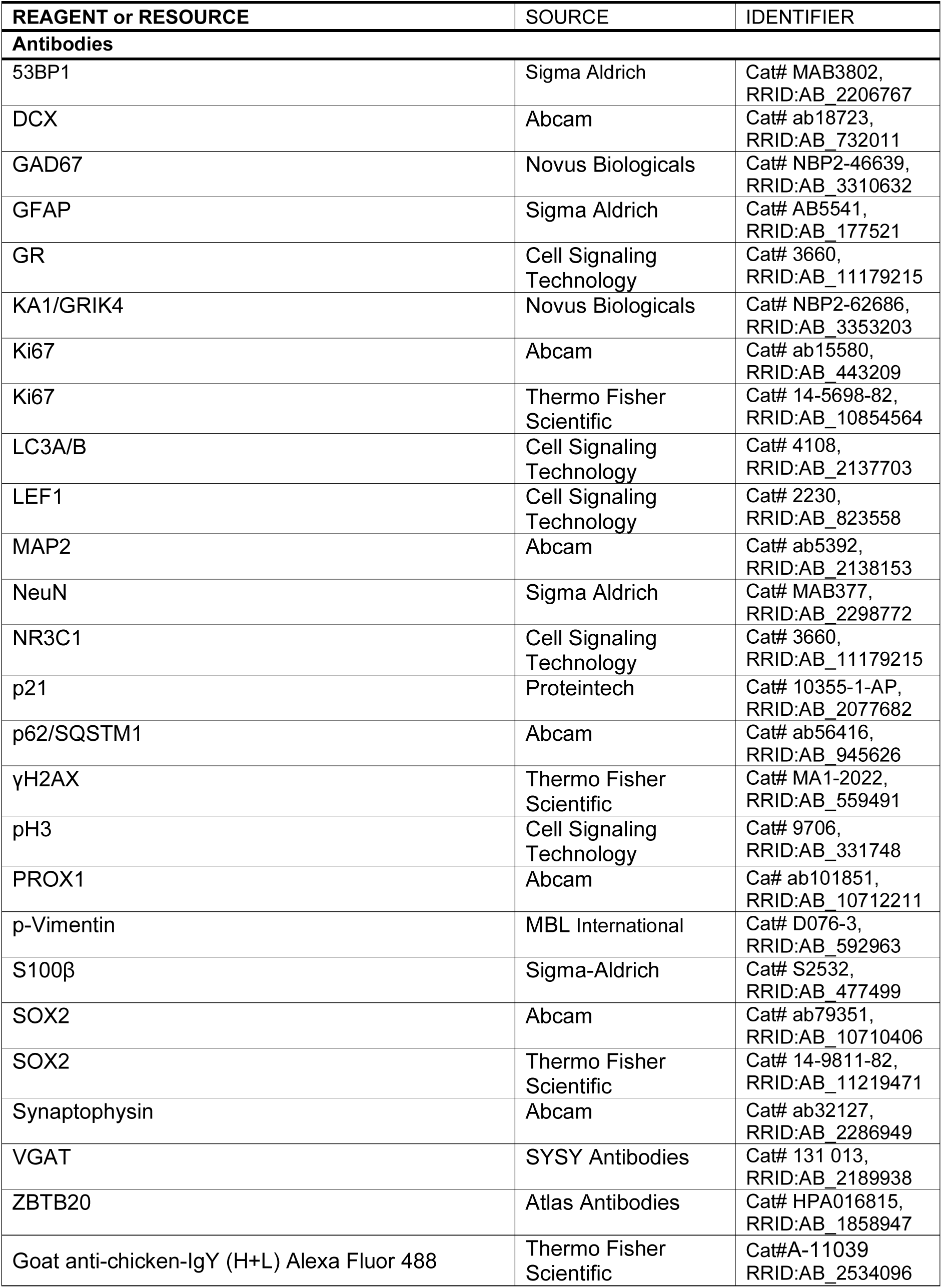
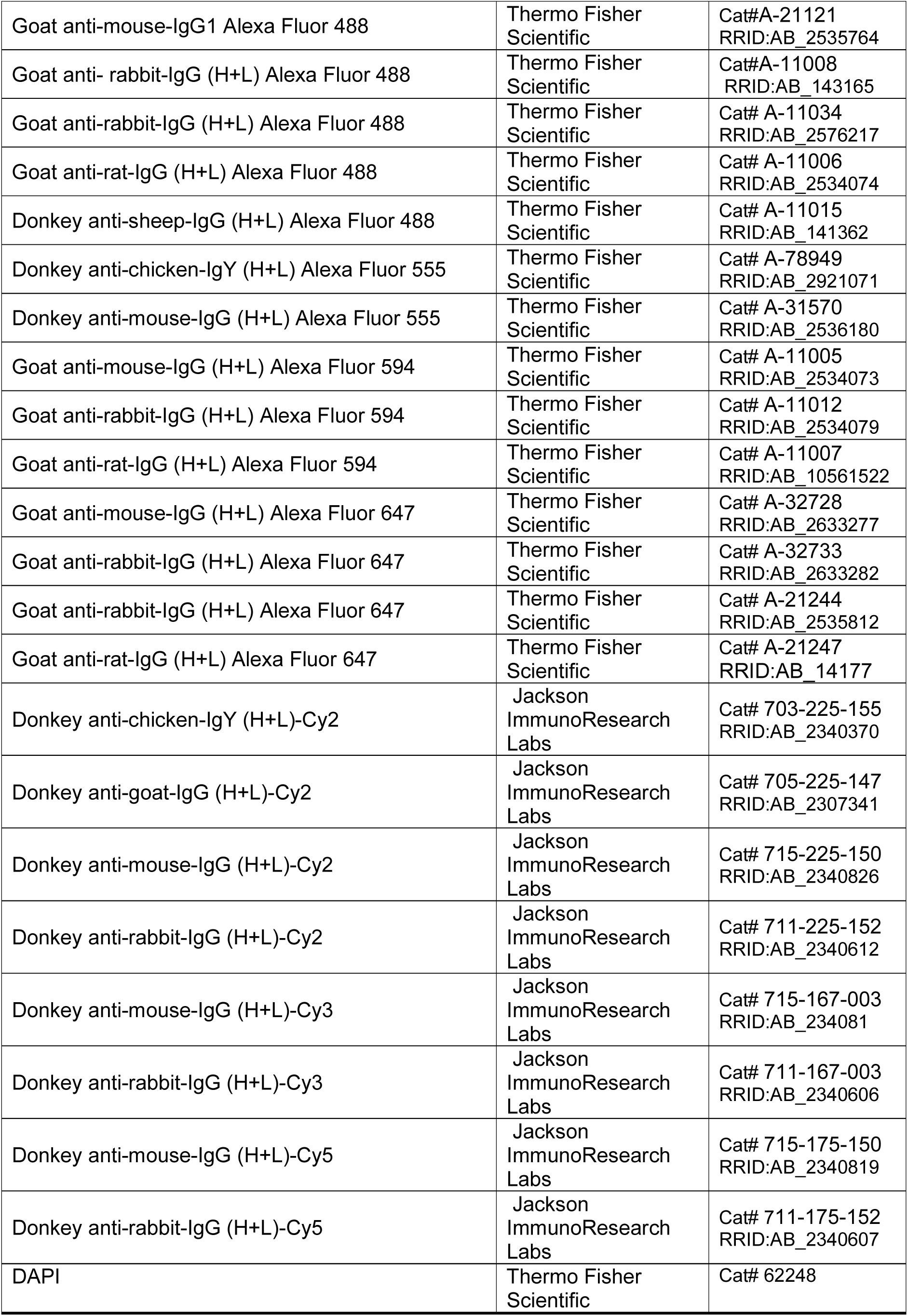
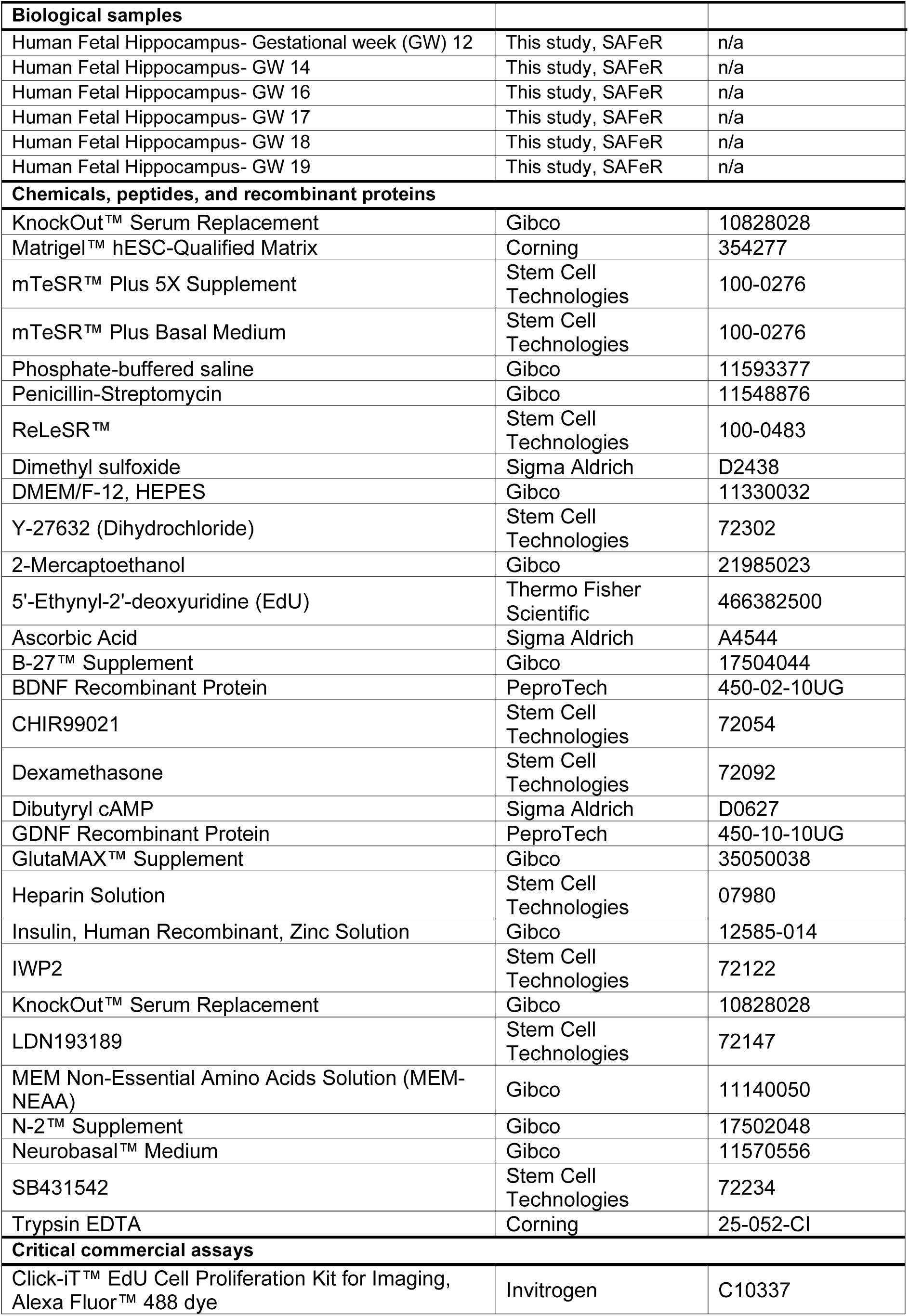
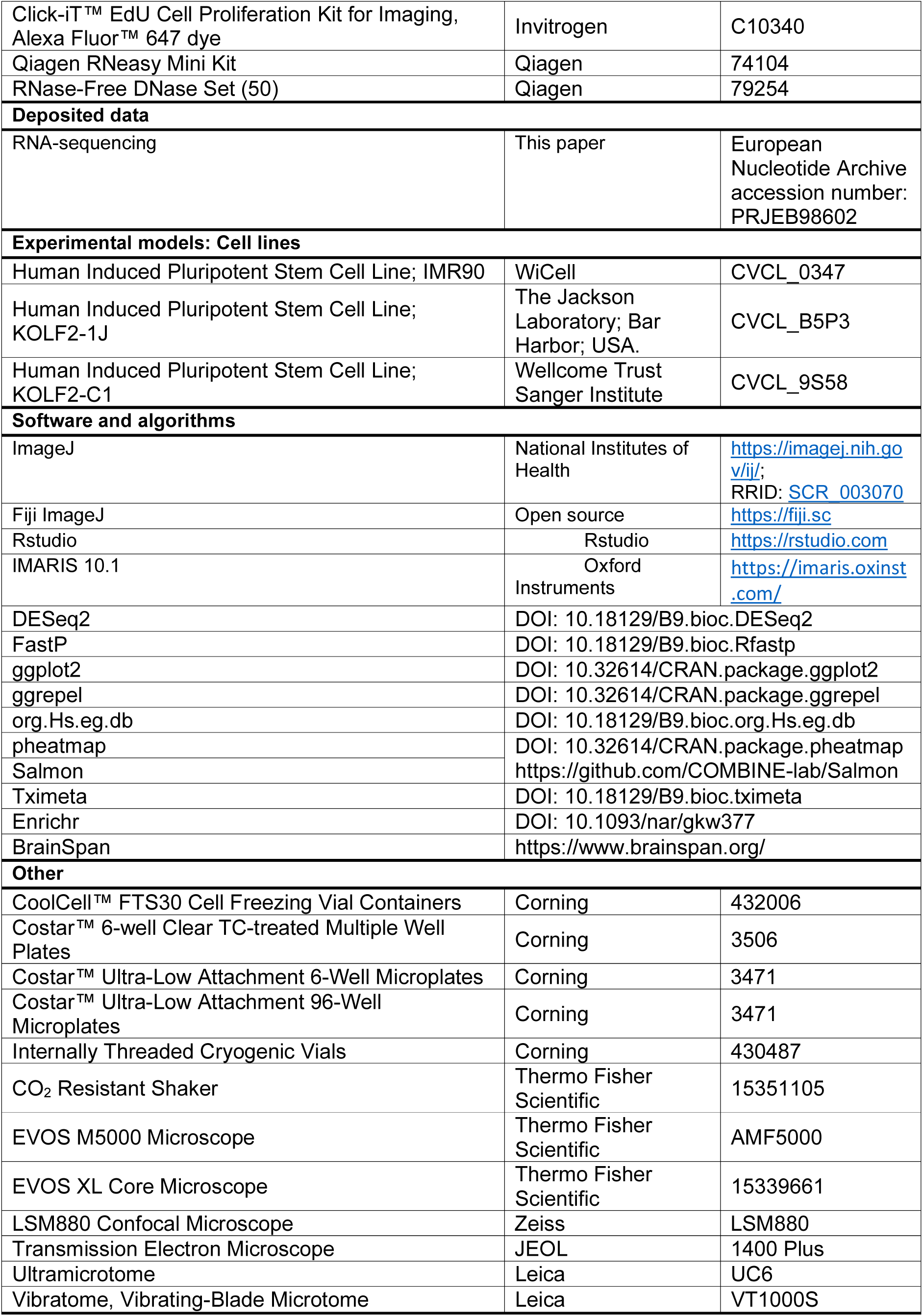

### Experimental model and study participant details

#### Cell Lines Used in this Study

iPSCs, KOLF2-C1, KOLF2-1J, or IMR90, were previously characterized^69–71^. The KOLF2-1J subline was purchased from The Jackson Laboratory and was generated through CRISPR-Cas9 gene editing to correct an ARID2 mutation present in the parental KOLF2-C1 line^69^. IMR90 cells were obtained from WiCell at passage P34 with a verified normal karyotype and were contamination-free.

#### Human Tissue Sample Collection

Fetal tissues from elective terminations of normally progressing pregnancies were obtained through the Scottish Advanced Fetal Research (SAFeR) study (NCT04613583)^72^. Collection was approved by one of the 12 Scottish National Health Service Research Ethics Committees (REC 15/NS/0123) and conducted in accordance with the Declaration of Helsinki. Recruitment was carried out by NHS Grampian research nurses, independent of the research team, who obtained written informed consent from women seeking elective medical termination of pregnancy. Eligible participants were women over 16 years of age, fluent in English, with pregnancies confirmed as normal by ultrasound, spanning gestational ages 7-20 weeks. Cases involving grossly abnormal fetuses were excluded, and women experiencing significant distress were not approached. Termination was performed using RU-486 (Mifepristone, 200 mg), followed by prostaglandin-induced delivery, as described previously^73^. Gestational age was confirmed by both ultrasound and fetal foot length measurement. Maternal and fetal demographic and clinical data were recorded. For sample collection, fetuses were immediately transported to the laboratory, typically intact, and subsequently weighed, measured for crown-rump length, and sexed based on external morphology. PCR analysis targeting sex chromosome-specific genes (ZFX and SRY) or absence of the Y chromosome was used to confirm sex assignment. All fetal hippocampal tissue samples analyzed in this study were derived from male fetuses. All samples were anonymized and identified only by unique study codes.

#### Method details

##### iPSC Culture

Human iPSCs were maintained on Matrigel-coated 6-well plates in mTeSR Plus medium (STEMCELL Technologies) at 37°C, 5% CO₂. For thawing, cryopreserved iPSCs were rapidly thawed, centrifuged at 300 × g for 3 min, and plated at ∼50,000 cells/well in medium supplemented with 10 μM Y-27632 (ROCK inhibitor). Medium was replaced daily. Cells were passaged at ∼80% confluence using ReLeSR and replated at 10,000-30,000 cells/well on freshly coated plates with Y-27632.

##### Generation of Hippocampal Organoids

The generation of hippocampal organoids was adapted from published cerebral and hippocampal protocols^28,29,32^ with modifications to optimize reproducibility and hippocampal specification. Once iPSCs reached approximately 80% confluence, they were used to generate embryoid bodies (EBs). The day of EB generation was defined as day 0 (d0). At d0, iPSCs were harvested into a 15 ml tube and centrifuged at 300 × g for 3 minutes. The supernatant was aspirated, and the pellet was resuspended in mTeSR Plus containing 10 μM Y-27632 ROCK inhibitor. This suspension was further diluted in mTeSR Plus with 10 μM Y-27632 to a final density of 2 × 10⁵ cells/ml. The cell suspension was then added to an ultra-low attachment 96-well U-bottom plate, typically at 5 × 10⁴ cells per well (250 μl/well). Any deviations from this starting density are noted in the respective Results sections. Plates were centrifuged at 300 × g for 5 minutes at 4°C and returned to the incubator.

At d2, 48 hours after EB formation, each EBs were collected from U-bottom well using a P1000 pipette with the tip cut. The EBs were pooled into a 15 ml tube and washed with DMEM/F12 and neural induction medium: H1 (for P1) or HO1 (for P2, P3, and P4). EBs were then evenly distributed into wells of a 6-well plate. Once the EBs had settled, the medium was aspirated and replaced with 3 ml of neural induction medium supplemented with 10 μM Y-27632 per well. Plates were incubated at 37°C on an orbital shaker (120 rpm) for 48 hours, after which the medium was replaced with H1 or HO1 lacking Y-27632.

Organoids were maintained on a shaker and media was changed every 48 hours, typically 3-4ml of media per well (see below for media compositions for each time point).

In P2, the base media and small molecule composition were altered. Additionally, gradual transitions were introduced between each base media change. Specifically, between d20 and d26, the media was progressively shifted from HO3 to HO4, as detailed below.

In P3, EBs were dissociated at d10. Briefly, EBs were collected into a 15 ml tube and washed with room temperature PBS. After removal of PBS, ∼500 μl of pre-warmed trypsin was added, and EBs were incubated at 37°C for 5 minutes. Mechanical dissociation was performed by gentle pipetting. HO2 medium was added, and the dissociated cells were centrifuged at 300 × g for 5 minutes. The supernatant was discarded, and the pellet was resuspended in HO3 medium supplemented with 10 μM Y-27632 ROCK inhibitor. Cells were seeded into a 96-well U-bottom plate at 250 μl per well, with the goal of reaggregating the same number of EBs as originally formed. Plates were centrifuged at 300 × g for 5 minutes to promote reaggregation and then returned to the 37°C incubator. Reaggregated EBs were maintained in the 96-well U-bottom plate between d10 and d20. Approximately 200 μl of HO3 medium was replaced with fresh HO3 medium every 48 hours until d18. On d20, reaggregated EBs were resuspended following the procedure above, washed with HO3 medium, and transferred into 6-well plates. Once settled, the medium was aspirated and replaced with 3 ml per well of HO3:HO4 (3:1) medium. Subsequent media changes were performed as described for P2.

In P4, on d10, EBs were transferred into a 1.5 ml Eppendorf tube using a cut P1000 pipette tip, the supernatant was removed, and EBs were washed twice with 1 ml HO3 medium. After the final wash, residual HO3 medium was removed, and 67 μl of fresh HO3 medium was added. To embed the EBs, 100 μl of thawed Matrigel was gently mixed with the 67 μl EB suspension, maintaining a 2:3 ratio of HO3 medium to Matrigel. The EB/HO3/Matrigel mixture was transferred to the center of an ultra-low attachment 6-well plate and spread into a thin, uniform layer to form a “Matrigel cookie” using a cut P200 pipette tip. If needed, EBs were repositioned with a P10 pipette tip to ensure even distribution and complete encapsulation. The plate was incubated at 37°C for 30 minutes to allow the Matrigel to solidify, after which 3 ml of pre-warmed HO3 medium was carefully added along the wall of each well to avoid disturbing the cookie. Plates were returned to the incubator and maintained without shaking. Medium was replaced every 48 hours with pre-warmed HO3.

On d20, organoids were gently extracted from the Matrigel cookie using a 5 ml serological pipette, transferred into a 15 ml tube, and washed three times with 5 ml room-temperature PBS. Organoids were then resuspended in 3 ml HO3:HO4 (3:1) medium and transferred to a standard 6-well plate placed on a shaker at 120 rpm in the incubator. From d22 onwards, media changes followed the same schedule as P2.

Media compositions

For P1,

- H1 (Neural induction media, day2-5): 20% KOSR, supplemented by 1XGlutamax, 1X MEM-NEAA, 0.1mM 2-Mecarptoethanol, 1 µM ldn193189, 5 µM SB431542, 0.2% Heparin Solution (2 µg/ml), and 1x penicillin/streptomycin in DMEM/F12 medium
- Day 6-7, Mixture of H1 and H2, H1:H2 = 1:1
- H2 (Hippocampal specification media, day8-15): 1X N2, 1X Glutamax, 1X MEM-NEAA, 0.1mM 2-Mercaptoethanol, 1x penicillin/streptomycin, 1 µM SB431542, 3 µM CHIR99021, and 20ng/ml BMP7 in DMEM/F12 medium
- H3 (Neurogenesis media, day 16-44): 1X N2, 1X B27, 1X Glutamax, 1X MEM-NEAA, 0.1mM 2-Mercaptoethanol, 1x penicillin/streptomycin and 2.5µg/ml Insulin in 50% DMEM/F12 and 50% Neurobasal medium
- H4 (Neural maturation media, day 45 onward): 1X B27, 1X Glutamax, 1X MEM-NEAA, 0.1mM 2-Mercaptoethanol, 1x penicillin/streptomycin, 0.05mM dibutyryl cAMP, 0.2mM Ascorbic acid, 20ng/ml BDNF and 20ng/ml GDNF in Neurobasal medium

For P2, P3, and P4,

- HO1 (neural induction media, day 2-5): 20% KOSR, supplemented by 1XGlutamax, 1X MEM-NEAA, 0.1mM 2-Mecarptoethanol, 1 µM LDN193189, 5 µM SB431542,

0.2% Heparin Solution (2 µg/ml), 1 µM IWP2, and 1x penicillin/streptomycin in DMEM/F12 medium

- Day 6-7, Mixture of HO1 and HO2; HO1:HO2 = 1:1
- HO2 (Hippocampal specification media 1, day8-9): 1X N2, 1X Glutamax, 1X MEM-NEAA, 0.1mM 2-Mercaptoethanol, 1x penicillin/streptomycin, 3 µM CHIR99021, and 20ng/ml BMP7 in DMEM/F12 medium
- HO3 (Hippocampal specification media 2, day 10-19): 1X N2, 1XB27, 1X Glutamax, 1X MEM-NEAA, 0.1mM 2-Mercaptoethanol, 1x penicillin/streptomycin, 1.5 µM CHIR99021, 10ng/ml BMP7, 0.5µg/ml Insulin in DMEM/F12 medium
- Day 20-25, Mixture of HO3 and HO4; day 20-21, HO3:HO4 = 3:1; d22-23, HO3:HO4 = 1:1; day 24-25, HO3:HO3 = 1:3
- HO4 (Differentiation and maturation media, day 26 onward):1X N2, 1X B27, 1X Glutamax, 1X MEM-NEAA, 0.1mM 2-Mercaptoethanol, 1x penicillin/streptomycin, 0.5µg/ml Insulin 0.05mM dibutyryl cAMP, 0.2mM Ascorbic acid, 10ng/ml BDNF in 50% DMEM/F12 and 50% Neurobasal medium
- HO5 (Neural maturation media, day 70 onward): 1X N2, 1X B27, 1X Glutamax, 1X MEM-NEAA, 0.1mM 2-Mercaptoethanol, 1x penicillin/streptomycin, 0.5µg/ml Insulin 0.05mM dibutyryl cAMP, 0.2mM Ascorbic acid, 10ng/ml BDNF in Neurobasal medium

##### Histology and Immunostaining

Hippocampal organoids were collected in 1.5 ml Eppendorf tubes, typically three per experimental timepoint. Organoids were washed with PBS and fixed in 4% paraformaldehyde (PFA) under a safety hood for 30-45 minutes at room temperature, depending on organoid size. Fixed samples were washed with PBS, cryoprotected in 30% sucrose, and stored at 4°C for a minimum of 24 hours before embedding in OCT freezing medium. Organoids were placed in OCT in cryomolds, frozen on dry ice, and stored at −80°C.

Human fetal hippocampus samples were placed in a 6-well plate and fixed in 4% PFA overnight at 4°C. The following morning, PFA was removed and replaced with 15% sucrose for 12 hours at 4°C, then with 30% sucrose for at least 24 hours. Samples were further dissected if required, embedded in OCT, frozen on dry ice, and stored at −80°C.

Frozen organoid and fetal tissue samples were sectioned at −20°C using a Leica CM1850 cryostat. Samples were embedded in OCT, and serial sections of 20 μm thickness were collected on Epredia SuperFrost Plus Adhesion Slides, air-dried overnight at room temperature, and stored at −20°C.

Sections were equilibrated to room temperature prior to staining, washed, and permeabilized with 0.05% Triton X-100 in 1X Tris-buffered saline (TBS+) for 5 minutes, repeated three times. For most antibodies, antigen retrieval was performed by incubating sections in 1X DAKO Target Retrieval solution at 85°C for 20 minutes in a plastic Coplin jar, followed by cooling to room temperature and three washes in TBS+. Blocking was carried out for 1 hour at room temperature in 5% bovine serum albumin (BSA) with 5% normal donkey or goat serum in TBS+, depending on the host species of the secondary antibodies.

Primary antibodies were diluted in antibody diluent (TBS+ containing 1.5% BSA and 1.5% normal donkey or goat serum) and incubated overnight at 4°C in a humidified chamber. Sections were washed three times in TBS+ before incubation with species-specific secondary antibodies for 2 hours at room temperature. Secondary antibodies were diluted in antibody diluent at 1:600 (Cy-conjugated) or 1:500 (Alexa Fluor-conjugated), with DAPI added at 1:10,000. Sections were washed three times with TBS+ and coverslipped using mounting medium prepared in-house (24% (w/v) glycerol, 9.6% (w/v) PVA,48% (v/v) 0.2 M Tris-HCl buffer, pH 8.0-8.5, and 24% (v/v) double-distilled water).

#### Senescence-Associated β-Galactosidase Staining

Senescence-associated β-galactosidase staining was performed according to the manufacturer’s instructions. Slides were washed with PBS and incubated in 1× Fixative Solution for 15 min at room temperature, followed by two PBS washes. A 20 mg/ml X-gal stock solution was prepared by dissolving 20 mg X-gal in 1 ml DMF. The working solution was prepared by mixing 50 μl X-gal stock with 930 μl 1× Staining Solution, 10 μl 100× Solution A, and 10 μl 100× Solution B. The pH was adjusted to 6.0 to ensure staining specificity. Slides were covered with staining solution, placed in a humidified chamber to prevent evaporation, and incubated at 37°C for ∼16 h until a blue signal developed.

#### EdU Pulse-Chase Experiments

For 5-ethynyl-2’-deoxyuridine (EdU) labeling of proliferating cells, hippocampal organoids were pulsed with 10 μM EdU in fresh culture medium at 37°C for 2 hours. After the pulse, the EdU-containing medium was removed, and organoids were washed once with PBS to eliminate residual EdU. Organoids were then replenished with fresh culture medium and returned to the incubator for the designated chase period. Media changes were performed every 48 hours during the chase, following routine culture conditions. Detection of EdU was performed after incubation with the secondary antibodies. Secondary antibodies were removed with three washes in TBS+, and the Click-iT EdU cell proliferation kit reaction cocktail was added following manufacturer’s instructions.

#### Dexamethasone Treatments

Dexamethasone was administered to hippocampal organoids on days 44, 46, and 48 of culture. A 100 μM dexamethasone stock was prepared in DMSO and diluted in HO4 medium to a final working concentration of 100 nM. Vehicle controls received an equivalent volume of DMSO.

The timing and concentration of treatment were adapted from Krontira et al.^74^. The selected concentration of 100 nM was designed to model antenatal corticosteroid therapy commonly used during pregnancies at risk of preterm birth. Clinical guidelines recommend intramuscular injection of four 6 mg doses of dexamethasone every 12 hours^75^. A single 6 mg dose reaches a peak plasma concentration (Cmax) of ∼65-95 ng/μl at ∼3 hours (Tmax), corresponding to ∼162-245 nM. The maternal-to-fetal steroid hormone ratio is ∼0.4 and increases in the days following treatment. Thus, after a single 6 mg dose, ∼65-98 nM dexamethasone would be expected to reach the fetus within 3 hours^74,76^.

Steroid hormones, including dexamethasone, are lipophilic and show strong binding to plastic. In human cortical organoid studies, 100 nM of applied androgens resulted in measured concentrations of ∼16 nM^74,77^. This reduction highlights the discrepancy between nominal and effective concentrations in vitro. Therefore, administering 100 nM dexamethasone in organoid cultures was considered physiologically relevant to fetal exposure levels observed in vivo during antenatal therapy.

In clinical practice, dexamethasone is administered as four doses across two days. With a biological half-life of 36-72 hours, fetal exposure is estimated to last 5-7 days, peaking within 2-3 days and gradually declining thereafter^74^. To reflect this pharmacokinetic profile, organoids were treated over a 6-day window.

#### Image Acquisition

Brightfield images were acquired using an EVOS XL Core microscope with 4× and 10× objectives. The phase ring was set to 20×/40× for images acquired with the 4× objective and to 4×/10× for images acquired with the 10× objective.

Fluorescent images were captured using an EVOS M5000 or a Zeiss LSM880 confocal microscope. For the EVOS M5000, the phase ring was set to 4×/10× for all samples. For quantification, at least three images were acquired per organoid section, with fields of view selected using DAPI staining to avoid bias. Confocal imaging was performed on a Zeiss LSM880 using 20×, 40× (oil immersion), and 63× (oil immersion) objectives. Laser power and detector gain were kept constant across samples for all intensity-based analyses.

For Transmission Electron Microscopy, samples were fixed in 2.5% glutaraldehyde and 4% PFA in 0.1 M sodium cacodylate buffer at 4°C for 24 h, then stored in 0.1 M sodium cacodylate buffer. Processing was performed using a Pelco Biowave Plus system. Tissues were washed in 0.1 M sodium cacodylate buffer (3 × 40 s, 150 W, 20 Hg vacuum) and post-fixed in 1% osmium tetroxide/1.5% potassium ferrocyanide in water (five cycles of vacuum on/off, 2 min each at 100 W, 20 Hg). After rinsing in distilled water (150 W, 20 Hg), samples were dehydrated through graded ethanol (35%, 50%, 70%, 95%) for 40 s each, followed by three washes in 100% acetone (40 s each, 150 W, 20 Hg). Tissues were infiltrated with graded acetone:Spurr’s resin (3:1, 1:1, 1:3) at 350 W, 20 Hg for 3 min, embedded in Spurr’s resin at 60°C for 48 h, and sectioned at 90 nm using a Leica UC6 ultramicrotome with a diamond knife. Sections were collected on formvar/carbon-coated copper grids, imaged on a JEOL 1400 Plus transmission electron microscope, and captured with an AMT UltraVUE camera. All TEM sample processing was performed by the Microscopy and Histology Core Facility, University of Aberdeen.

#### Image Analysis

All image analysis was performed using Fiji (ImageJ)^78^. To minimize bias, nuclear markers, including DAPI, were quantified using a standardized pipeline. Images were converted to binary, and nuclei were segmented using the Auto Local Threshold tool. Where necessary, the Watershed tool was applied to separate adjacent nuclei. The Analyze Particles function was then used to quantify nuclei and extract descriptors such as size. For analyses limited to specific regions of interest (ROIs), such as SOX2^+^ NSC layers, ROIs were generated from masks of the relevant marker, restricting quantification to nuclei within those defined regions.

Due to high variability in expression levels of PROX1 and ZBTB20 across developmental timepoints, these markers were quantified manually. Marker fluorescence channels were overlaid with DAPI nuclear staining, and the Channels Tool was used to toggle between channels to confirm colocalization. Nuclei were considered positive when the marker signal overlapped with a DAPI-stained nucleus. Positive nuclei were annotated with the Cell Counter plugin, ensuring each nucleus was counted once across Z-stacks.

Organoid morphology was analyzed from brightfield images, ensuring each organoid was counted only once. Morphological parameters, including area, perimeter, Feret’s diameter, and circularity, were measured. Circularity was calculated as 4π(area/perimeter²), with 1.0 indicating a perfect circle. Feret’s diameter was defined as the maximum distance between two boundary points. Images were binarized, artefacts removed using the Remove Outliers tool, and edges detected with Find Edges. Manual refinement with the Paintbrush tool was applied when necessary to isolate individual organoids.

Outgrowths were counted manually from 4× brightfield images. Outgrowths were defined as solid-edged, translucent extensions, either attached or detached from the organoid.

For fluorescence localization, SOX2 and DCX intensity profiles were obtained using the Plot Profile command. A line was drawn from the center of an NSC layer (or the organoid core, where specified) to the edge, and signal intensity was measured along this line. Curves were generated in GraphPad Prism (v8.4.3) using LOWESS smoothing.

The number of NSC layers was counted manually. NSC layer area was measured by converting images to binary and selecting the layers with the Wand tool. Thickness was measured by drawing a line from the center of the SOX2^+^ NSC layer to its periphery. For immature neurons, thickness was measured from the edge of the NSC layer to the organoid surface using DCX staining.

The total area covered by specific markers was quantified by converting images to binary, generating a mask with the Auto Local Threshold tool, and measuring the area of positive signal. Fluorescence intensity was quantified using the same pipeline, with masks generated from the relevant marker or from SOX2 for NSC-specific analyses. Integrated density was used when marker-positive area differed between conditions, as it reflects both signal intensity and size. Mean gray value was used when the marker area was consistent across groups, enabling relative comparisons of intensity per unit area. This ensured that signal differences reflected true expression changes rather than variation in localization or cell number.

#### Human Tissue Dissection

Whole fetal brains were transferred to ice-cold Hanks’ Balanced Salt Solution (HBSS). Brains were then placed in sterile 90 mm petri dishes filled with HBSS and dissected on ice. Key anatomical landmarks, including the Sylvian fissure, central sulcus, cerebellum, and brainstem, were used to orient and guide dissection. The hemispheres were separated by a sharp midline incision through the brainstem. To identify the hippocampus, medial surface landmarks such as the developing corpus callosum and fornix were located. The hippocampus was dissected in a septal-to-temporal direction, and excess cortical tissue was carefully removed. Dissected samples were either fixed for immunohistochemistry or processed immediately for RNA extraction.

#### RNA Sequencing

Total RNA was extracted from hippocampal organoids or human fetal hippocampal tissue using the *RNeasy Mini Kit* (Qiagen) according to the manufacturer’s protocol, including on-column DNase I digestion to remove genomic DNA contamination. RNA quality and integrity were assessed using an *Agilent Bioanalyzer 2100* (Agilent Technologies), and samples with RNA integrity number (RIN) ≥ 7 were used for sequencing. RNA concentration was quantified using a *Qubit Fluorometer* (Thermo Fisher Scientific). RNA sequencing of hippocampal organoid development in comparison to fetal hippocampi was done by a NextSeq 500 machine at the Centre for Genome Enabled Biology and Medicine (CGEBM), University of Aberdeen. RNA sequencing for dexamethasone-treated hippocampal organoids was processed by stranded RNA sequencing performed on Illumina Novoseq X Plus by Novogene.

#### RNA-Sequencing Data Processing and Analysis

RNA-sequencing analysis was performed as previously described^79^. Raw FASTQ files were concatenated by sample and processed with FASTP, which performs quality profiling, adapter trimming, and removal of low-quality reads^80,81^.

Cleaned reads were mapped to the human reference genome (GRCh38) and quantified at the transcript level using Salmon v1.10.2 in mapping-based mode^82^. A Salmon index was generated from Ensembl GRCh38 cDNA FASTA files, and transcript-level quantification was obtained.

Quantification files were imported into R v4.5.0 using the *tximeta* Bioconductor package (v3.19), generating a *SummarizedExperiment* object with gene-level count matrices and annotation metadata^83^. Ensembl gene IDs were converted to gene symbols using *org.Hs.eg.db*. Differential expression analysis was conducted with DESeq2^84^, excluding genes with fewer than two non-zero counts across all samples. Genes with |log2 fold change| ≥ 0.4 and an adjusted *p* < 0.05 were considered differentially expressed. Results were exported as .csv files, and visualizations were generated using *ggplot2*/*ggrepel* (volcano plots) and *pheatmap* (heatmaps).

Functional enrichment of differentially expressed genes (DEGs) was performed using Enrichr^85^. Developmental relevance was assessed by integrating data from the BrainSpan atlas (https://www.brainspan.org/), filtering for genes overlapping with the organoid dataset, and calculating Spearman’s correlations between organoids and human brain regions, including the hippocampus, across developmental stages.

For organoid characterization, curated marker gene sets from Ciarpella et al. were used, including markers for neural stem/progenitor cells^86^, mature neurons^86–88^, astrocytes^88,89^, and excitatory/inhibitory synapses^90^. Autophagy-associated gene sets linked to NSC quiescence were obtained from Calatayud-Baselga et al.^59^, while active-to-quiescent NSC transition modules were derived from Jimenez-Cyrus et al.^58^.

#### Vibratome Sectioning

Organoids were sectioned after day 90 of culture when required for electrophysiology or calcium imaging. Organoids were embedded in 3% agarose in custom molds and sectioned in DMEM/F12 using a Leica VT1000S vibratome (Leica Microsystems) at 0.1 mm/second, 80 Hz frequency, and 1.00 mm amplitude. Sections were cut at 300 μm thickness, collected with a cut-Pasteur pipette, and transferred to HO5 medium. Slices were maintained under standard culture conditions on an orbital shaker at 120 rpm.

#### Calcium Imaging

Calcium imaging was performed using Fluo-4 AM. Organoids were incubated for 1 hour at 37°C in a 1:1 solution of 10 μM Fluo-4 AM and Pluronic prepared in HO3 or HO4 medium, depending on organoid age. Prior to imaging, organoids were washed with HBSS without calcium and magnesium, then incubated for 5 minutes at room temperature in HBSS containing calcium and magnesium.

Immediately before imaging, most of the HBSS was removed to prevent organoid drifting, leaving a small residual volume to avoid sample drying. Fluorescence imaging was performed on an EVOS M5000 system equipped with a 10× objective and an onstage incubator set to 37°C, 5% CO₂, and 80% humidity. Time-lapse images were acquired continuously at the fastest possible acquisition rate of 10 minutes.

#### Calcium Imaging Analysis

Image analysis was performed using ImageJ. To correct for organoid drift, image stacks were aligned with the Linear Stack Alignment with SIFT plugin^91^. A maximum intensity projection of each stack was generated to identify neurons, and ROIs were manually drawn around individual neurons. Fluorescence intensity within each ROI was measured across the entire time series.

Fluorescence changes were quantified as ΔF/F = (Fstim - Fbasal) / Fbasal, where Fstim represents the fluorescence intensity of a neuron at a given time point, and Fbasal represents baseline fluorescence measured from cell-free regions of the organoid. Neurons were classified as active if calcium transients were detected at least once during the imaging period. Heatmaps were generated in R using the ComplexHeatmap package (v2.20.0)^92^.

#### Electrophysiology

Hippocampal organoids aged between day 70 and day 160 were used for whole-cell patch-clamp recordings. On the day of recording, organoids were transferred to a recording chamber, secured with a weighted harp, and continuously perfused with oxygenated bath solution containing 125 mM NaCl, 3 mM KCl, 1.2 mM NaH2PO4, 26 mM NaHCO3, 1 mM MgCl2, 2 mM CaCl2, and 25 mM glucose, bubbled with 95% O₂ and 5% CO₂. Individual neurons were visualized using an upright microscope equipped with infrared illumination, and all recordings were performed at room temperature.

Recording pipettes were pulled from borosilicate glass capillaries to a resistance of 5.3-7.5 MΩ and filled with intracellular solution containing 130 mM K-gluconate, 10 mM HEPES, 0.1 mM EGTA, 5 mM NaCl, 4 mM MgATP, 0.3 mM Na₃GTP, and 2 mM pyruvic acid.

Whole-cell recordings were obtained using Axon Instruments hardware (MultiClamp 700B, Digidata 1550) and pCLAMP 10.7 software. Data were low pass filtered at 2 kHz and sampled at 10 kHz. Input resistance, rheobase, and action potential threshold were assessed in current-clamp mode.

For input resistance, neurons were held near −50 mV by current injection, and a series of hyperpolarizing current steps (250 ms, 5 pA increments) were applied until the membrane potential reached approximately −100 mV. Input resistance was calculated as the slope of the voltage-current relationship measured at the end of the hyperpolarizing steps.

For rheobase (minimum current required to elicit an action potential) and action potential threshold, neurons were held near −50 mV and short (3 ms) depolarizing current pulses were applied in 10 pA increments until an action potential was evoked.

#### Statistical Analysis and Data Visualization

Statistical analyses were performed using GraphPad Prism (v8.4.3). All statistical tests, sample sizes, and exact *p*-values are detailed in the figure legends. Data are presented as mean ± SEM, mean ± SD, or median ± interquartile range (IQR), as indicated. Statistical significance was defined as *p* ≤ 0.05, with thresholds reported as follows: *p* ≤ 0.05 (\**), p ≤ 0.01 (**), p ≤ 0.001 (\*\*\**), and *p* ≤ 0.0001 (****). Differences were considered not statistically significant when *p* > 0.05.

Data distributions were assessed for normality using the Shapiro-Wilk or Kolmogorov-Smirnov tests. For parametric data, comparisons were performed using unpaired (or paired, where appropriate) *t*-tests or one-way ANOVA with Tukey’s multiple comparisons test. For non-parametric data, Mann-Whitney or Kruskal-Wallis tests with Dunn’s multiple comparisons test were applied.

RNA-seq analyses were performed in the R statistical environment (v4.3.1, https://www.r-project.org/). Graphs and plots were generated using GraphPad Prism or R, with final edits completed in Adobe Illustrator 2025. All schematic figures were created in Adobe Illustrator 2025.

