## supplementary information for "A Human Hippocampal Organoid Model with Sustained Neural Stem Cells Reveals State Shifts Under Glucocorticoid Stress"

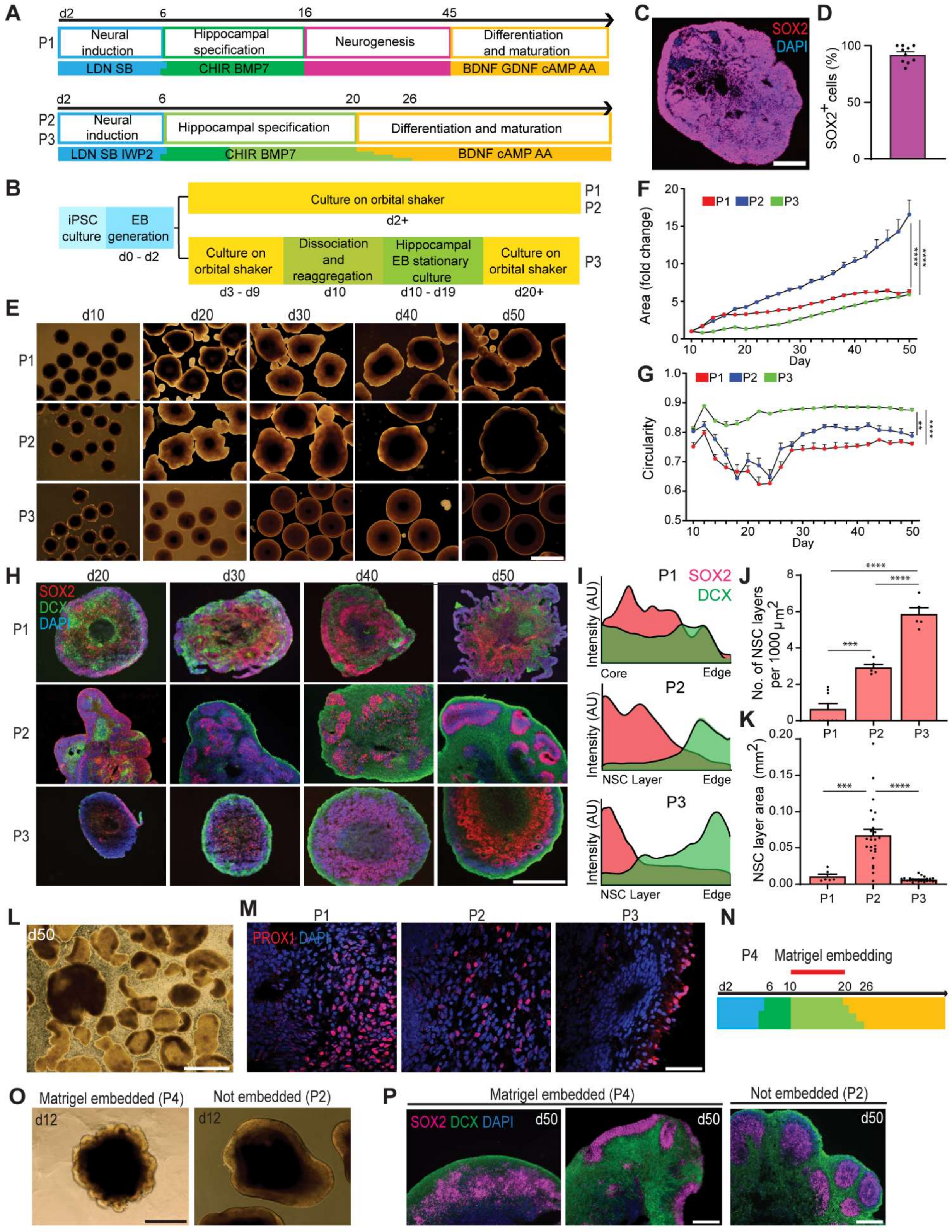
A
P1
d2
6
16
45
Neural induction
Hippocampal specification
Neurogenesis
Differentiation and maturation
LDN SB
CHIR BMP7
BDNF GDNF cAMP AA
P2
P3
d2
6
20
26
Neural induction
Hippocampal specification
Differentiation and maturation
LDN SB IWP2
CHIR BMP7
BDNF cAMP AA
B
iPSC culture
EB generation
d0 - d2
Culture on orbital shaker
d2+
P1
P2
Culture on orbital shaker
d3 - d9
Dissociation and reaggregation
d10
Hippocampal EB stationary culture
d10 - d19
Culture on orbital shaker
d20+
P3
C
SOX2
DAPI
D
SOX2+ cells (%)
E
d10
d20
d30
d40
d50
P1
P2
P3
F
P1
P2
P3
Area (fold change)
Day
G
Circularity
Day
H
d20
d30
d40
d50
SOX2
DCX
DAPI
I
SOX2
DCX
Intensity (AU)
Core
Edge
NSC Layer
J
No. of NSC layers per 1000 μm2
K
NSC layer area (mm2)
L
d50
M
PROX1 DAPI
N
P4
Matrigel embedding
d2
6
10
20
26
O
Matrigel embedded (P4)
Not embedded (P2)
d12
P
Matrigel embedded (P4)
Not embedded (P2)
SOX2 DCX DAPI
d50

**Figure S1. Comparison of protocols and optimisation strategies for hippocampal organoid generation. Related to Figure 1.**

**(A)** Schematic illustrating the four-phase media protocol (P1) and the three-phase protocol (P2, P3) used to develop and optimise hippocampal organoids (see STAR Methods for additional details). Gradual transitions between media phases are indicated by the stepwise colour changes.

**(B)** Schematic overview of the different protocols used to develop and optimise hippocampal organoids (see STAR Methods for additional details). To address the non-radial growth patterns that were observed in P1 and occasionally in P2 during preliminary studies, a dissociation and reaggregation step was introduced in P3 at day (d) 10 of culture.

**(C-D)** Representative immunofluorescence image of neural stem cells (NSCs; SOX2, magenta) present in a P2-generated hippocampal organoid at d10 (**C**). The d10 timepoint was selected for dissociation and reaggregation in P3 based on observations showing that at this timepoint ~97% of the cells present in the organoid were SOX2$^+$ NSCs (**D**). It was hypothesised that perturbation of the cytoarchitecture in organoids predominantly composed of NSCs, prior to differentiation into DCX$^+$ immature neurons, would facilitate cellular rearrangement, thereby promoting radial growth. Scale bar = 200 µm. Data are presented as mean + SEM. n = 6 sections from 3 hippocampal organoids.

**(E)** Representative brightfield images of hippocampal organoids produced from P1, P2 and P3 at d10, d20, d30, d40 and d50. Scale bar = 500 µm.

**(F)** The growth of hippocampal organoids produced from P1, P2 and P3 over time. Organoids from all protocols were found to exhibit similar growth trends, with progressive increases in area as expected. However, P2 exhibited a higher growth rate, with a fold change in area of 16.57 ± 4.291 and overall area change of 6.78 ± 1.76 mm$^2$ by d50. In comparison, P1 and P3 resulted in a fold change in area of 6.33 ± 0.93 (overall area; 2.607 ± 0.38 mm$^2$) and 5.92 ± 0.75 (overall area; 2.42 ± 0.31 mm$^2$) respectively. Statistical differences between protocols determined by Two-Way ANOVA with Tukey's multiple comparisons test. Significance shown is for d50; **** $p < 0.0001$.

**(G)** The circularity of hippocampal organoids produced from P1, P2 and P3 over time. High circularity values are typically observed during early organoid culture, particularly in the EB phases. As neuroepithelial buds and lobes begin to form, organoid morphology becomes more complex and less spherical, leading to a decline in circularity. This reduction stabilises as these structures mature and are maintained over time. Therefore, the complexity of organoid morphology can be indicated by circularity measurements. P1 and P2 exhibited decreasing circularity between d10 and d28, reaching a minimum of 0.62 (d22) and 0.64 (d18) respectively, before stabilising at approximately 0.75 for P1 and 0.80 for P3 after d30. In contrast, P2 maintained relatively constant circularity throughout the culture period, ranging from a minimum of 0.82 at d16 to a maximum of 0.89 at d36. Statistical differences between protocols determined by Two-Way ANOVA with Tukey's multiple comparisons test. Significance shown is for d50; ** $p = 0.003$, **** $p < 0.0001$.

**(H)** Representative immunofluorescence images of hippocampal organoids produced from P1, P2 and P3 at different time points (d20 – d90), demonstrating the presence of NSCs (SOX2, magenta) and immature neurons (DCX, green) in both radial and non-radial growth patterns. Organoids generated using P1 were found to exhibit disorganised, non-radial cellular distribution, with SOX2$^+$ NSCs primarily localised at the organoid's periphery and DCX$^+$ immature neurons localised to the core of the organoid. Additionally, the absence of a well-defined border to the organoids was apparent. In contrast, organoids generated from both P2 and P3 were found to exhibit progressive cellular organisation during maturation. In P2, a lack of radial cellular organisation was apparent at d20 with $SOX2^+$ cells primarily located around the periphery of the organoid. However, $SOX2^+$ cells were also present in the centre, where $DCX^+$ cells were predominantly observed. However, by d30 there was evidence of cellular organisation with the formation of NSC layers and a switch in growth orientation, with the DCX+ cells now at the periphery of the organoid and SOX2+ NSC layers towards the centre of the organoid, thereby growing in a radial manner. This pattern was maintained at both d40 and d50. A similar trend was observed in P3 organoids, though with a delayed onset of organisation, with the lack of radial cellular organisation apparent at both d20 and d30. At d20, SOX2+ cells surround the periphery of the organoid and there was a lack of DCX+ cells. However, by d30, $SOX2^+$ cells are localised to the centre of the organoid, although in a disorganised manner, and $DCX^+$ cells surround the periphery of the organoid. By d40, distinct NSC layers become apparent with central localisation of $SOX2^+$ cells, and $DCX^+$ cells remain around the periphery of the organoid. These cellular organisation and radial growth patterns are maintained at d50. The delayed cellular organisation in P3 organoids is likely attributable to the dissociation and reaggregation step at d10 which may have reset or delayed their developmental progression. Scale bar = 500 μm.

**(I)** Fluorescent intensity analysis showing expression of SOX2 (red) and DCX (green) plotted against the distance from the core to the periphery of the organoid at d50 for P1, or from the middle of an NSC layer to the periphery of the organoid for P2 and P3 at d50, highlighting non-radial (P1) and radial (P2 and P3) growth trends.

**(J)** Quantification of the number of neural stem cell (NSC) layers at d50 in hippocampal organoids generated from P1, P2 and P3 at d50. P1-generated organoids exhibited minimal NSC layer formation, with an average of 0.53 NSC layers per 100 $\mu m^2$ in the organoids. In contrast, P2- and P3-generated organoids demonstrated significantly greater NSC layer formation, averaging 2.9 and 5.9 layers per 100 $\mu m^2$ in the organoids, respectively. Data are expressed as mean + SEM. Statistical differences between protocols were determined by ordinary one-way ANOVA followed by Tukey's multiple comparison test. n = 5 – 8 hippocampal organoids per protocol. *** $p = 0.0002$, **** $p < 0.0001$.

**(K)** Quantification of the area of NSC layers at d50 in hippocampal organoids generated from P1, P2 and P3. NSC layers present in P2-generated organoids were significantly larger than those present in P1- and P3-generated organoids, with an average area of 0.067 ± 0.04 $mm^2$ in P2 versus 0.01 ± 0.008 $mm^2$ in P1 and 0.006 ± 0.003 $mm^2$ in P3 at d50. Data are expressed as mean + SEM. Statistical differences between protocols were determined by ordinary one-way ANOVA followed by Tukey's multiple comparison test. n = 6 – 25 NSC layers from at least 3 hippocampal organoids per protocol. Statistical significance determined by One-Way ANOVA with Tukey's multiple comparisons test. *** $p = 0.0001$, **** $p < 0.0001$.

**(L)** Representative brightfield image of a d50 organoid that was displaying non-radial growth in P1. This disorganised cellular distribution prevented the long-term culture of organoids. The inward migration of neurons toward the centre of the organoid ultimately resulted in spatial constraints, leading to structural instability, fragmentation, and disintegration of the organoid. Scale bar = 1 mm

**(M)** Representative immunofluorescence images of dentate granule cells (PROX1, red) in d50 hippocampal organoids generated from P1, P2 and P3. Scale bar = 50 μm.

**(N)** Schematic illustrating the period of Matrigel embedding (d10 – d20) in P4 (see STAR Methods for additional details). Organoids were placed in Matrigel from d10 to d20, corresponding to the hippocampal specification stage of media provision

**(O)** Representative brightfield images of a hippocampal organoid embedded in Matrigel (P4, left) and a hippocampal organoid not embedded in Matrigel (P2, right) at d12, 48 hrs after Matrigel embedding. Prominent neuroepithelial bud-like structures were apparent in Matrigel-embedded organoids. Scale bar = 500 µm

**(P)** Representative immunofluorescence images of hippocampal organoids embedded in Matrigel (P4) displaying disorganised NSC layers (left), and non-radial growth (centre), compared with age- and batch-matched non-embedded controls (P2) displaying structured NSC layers and radial growth (right) at d50. These results suggest that Matrigel embedding negatively impacts hippocampal organoid growth and is therefore not recommended for optimal culture conditions. NSCs ($SOX2^+$) shown in magenta, immature neurons ($DCX^+$) shown in green. Scale bar = 200 µm

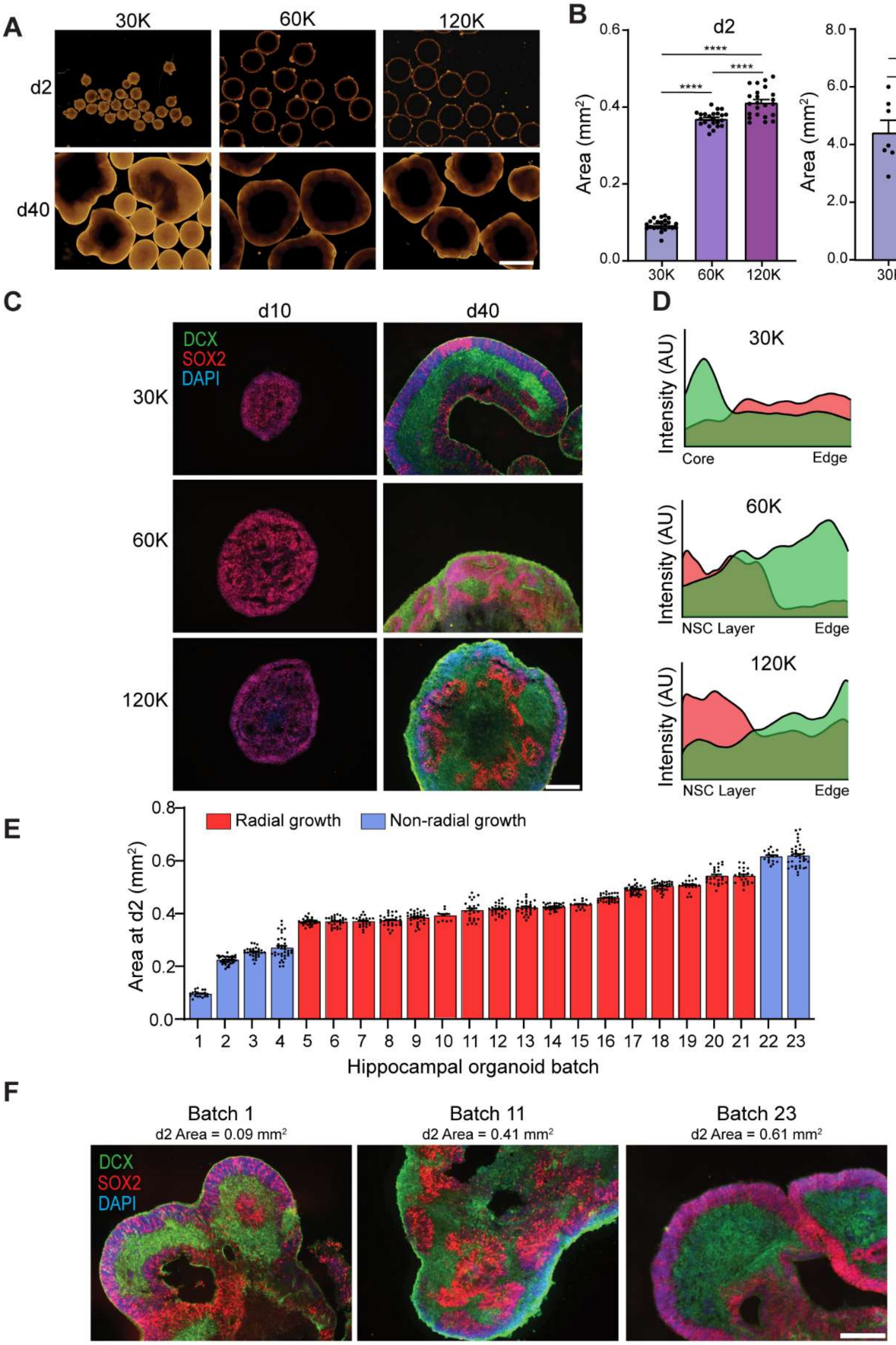

A
30K
60K
120K
d2
d40
B
d2
d40
Area (mm2)
30K
60K
120K
C
d10
d40
DCX
SOX2
DAPI
30K
60K
120K
D
30K
60K
120K
Intensity (AU)
Core
Edge
NSC Layer
E
Radial growth
Non-radial growth
Area at d2 (mm2)
Hippocampal organoid batch
F
Batch 1
d2 Area = 0.09 mm2
Batch 11
d2 Area = 0.41 mm2
Batch 23
d2 Area = 0.61 mm2
DCX
SOX2
DAPI

**Supplementary Figure 2. Initial EB size determines radial growth patterns in P2 hippocampal organoids. Related to Figure 1**

**(A)** Representative brightfield images of P2-generated hippocampal organoids generated from 30,000, 60,000 and 120,000 iPSCs at d2 and d40. The impact of initial EB size on the radial growth of the organoids was investigated by generating EBs from different numbers of iPSCs. Brightfield imaging revealed clear morphological differences between EBs generated from 30,000 iPSCs and those generated from 60,000 and 120,000 iPSCs, although minimal morphological differences were apparent between EBs generated from 60,0000 and 120,000 iPSCs. Scale bar = 1 mm.

(**B**) Area of P2 hippocampal organoids at d2 (left) and d40 (right) of culture. Morphology analysis of d2 EBs revealed a significant difference in area between all EB sizes. However, the average areas of the EBs generated from 60,000 and 120,000 iPSCs were closer than expected, being 0.37 $mm^2$ and 0.41 $mm^2$, respectively. Whereas EBs generated from 30,000 iPSCs had an average area of 0.09 mm2. Interestingly by d40, there were still significant differences between EBs generated from 60,000 iPSCs (average area = 2.83 $mm^2$) and 120,000 iPSCs (average area = 2.42 $mm^2$) compared to EBs generated from 30,000 iPSCs (average area = 4.42 $mm^2$) ($p < 0.0001$). However, no significant differences were found between EBs generated from 60,000 iPSCs and those generated from 120,000 iPSCs Data are presented as mean + SEM. n = 7 - 23 hippocampal organoids per condition. Statistical significance determined by One-Way ANOVA with Tukey's multiple comparisons test. **** $p < 0.0001$.

**(C)** Representative immunofluorescence images of P2 hippocampal organoids generated from 30,000, 60,000 and 120,000 iPSCs at d10 and d40, showing the neural stem cells (SOX2, red) and immature neurons (DCX, green). At d10 approximately 100% of cells in the EB were SOX2+ NSCs, regardless of starting EB size. However, by d40, organoids generated from 30,000 iPSCs displayed SOX2+ NSC layers at the periphery of the organoid, with DCX+ immature neurons localised towards the centre of the organoid, thereby growing in a non-radial manner. In contrast, organoids generated from both 60,000 iPSCs and 120,000 iPSCs displayed SOX2+ NSC layers within the centre of the organoid, with DCX+ immature neurons towards the edge of the organoid, growing in a radial manner. Scale bar = 1 mm.

**(D)** Fluorescent intensity analysis showing expression of SOX2 (red) and DCX (green) plotted against the distance from the core to the periphery of the organoid at d50 for P2 hippocampal organoids generated from 30,000 iPSCs, highlighting non-radial growth trends, or from the middle of a NSC layer to the periphery of the organoid for organoids generated from 60,000 and 120,000 iPSCs, highlighting radial growth trends, at d50.

**(E)** Quantification of d2 EB size of 23 batches of hippocampal organoids displaying non-radial (blue) and radial growth (red). The optimal d2 EB area to generate hippocampal organoids displaying radial growth is between 0.37 $mm^2$ and 0.54 $mm^2$. n = 9 – 38 hippocampal organoids per batch.

**(F)** Representative immunofluorescence images of P2 hippocampal organoids at d30 from different batches, showing the impact of d2 EB size on radial growth patterns. SOX2+ NSCs shown in red, DCX+ immature neurons shown in green. Scale bar = 200 μm.

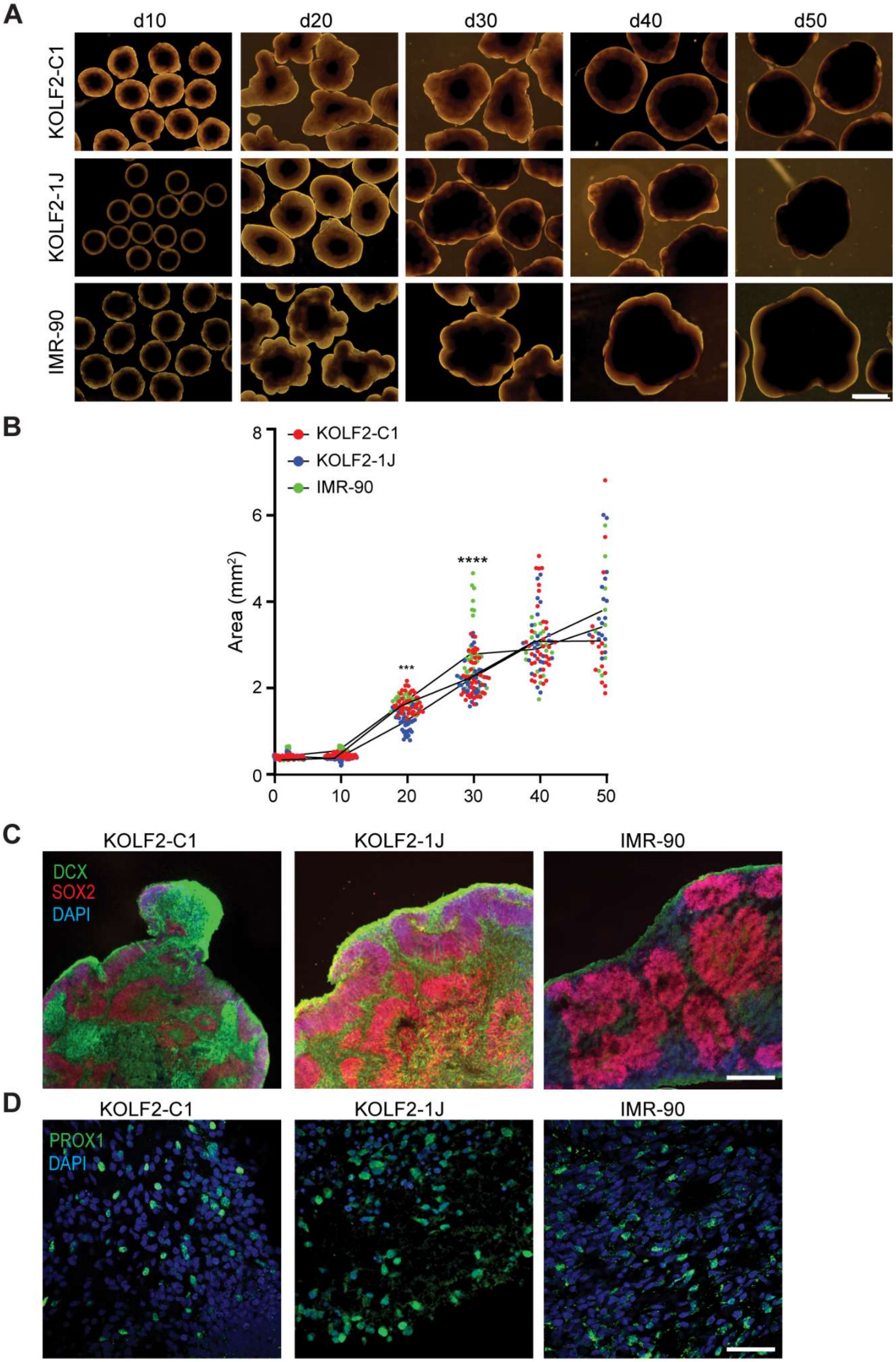
A
d10
d20
d30
d40
d50
KOLF2-C1
KOLF2-1J
IMR-90
B
KOLF2-C1
KOLF2-1J
IMR-90
Area (mm²)
***
****
0
2
4
6
8
0
10
20
30
40
50
C
KOLF2-C1
KOLF2-1J
IMR-90
DCX
SOX2
DAPI
D
KOLF2-C1
KOLF2-1J
IMR-90
PROX1
DAPI

**Supplementary Figure 3. Characterisation of hippocampal organoids generated from multiple iPSC lines. Related to Figure 1**

**(A)** Representative brightfield images show similar morphology of protocol (P) 2 hippocampal organoids generated from KOLF2-C1, KOLF2-1J and IMR90 iPSCs at day (d) 10, d20, d30, d40 and d50. Scale bar = 1 mm.

**(B)** Quantification of P2 organoid area from d2 to d50 demonstrates parallel growth trends across all three iPSC lines (KOLF2-C1, KOLF2-1J and IMR90 iPSCs). Statistically significant differences in size were observed at early timepoints (d20 - d30), but these differences were no longer present by d40 - d50. These differences may therefore be due to inherent line-to-line variability, which did not affect the organoid growth or maturation. n = 18 - 72 organoids per cell line per timepoint. 3 independent batches of organoids were analysed per cell line. Statistical significance determined by Two-Way ANOVA with Tukey's multiple comparisons test. d20: KOLF2-C1 vs. KOLF2-1J, *** $p = 0.0005$; IMR90 vs. KOLF2-1J, *** $p = 0.0008$; d30: IMR90 vs. KOLF2-C1, *** $p = 0.0001$; IMR90 vs. KOLF2-1J, **** $p < 0.0001$.

(**C**) Representative immunofluorescence images showing the neural stem cells (SOX2, magenta) and immature neurons (DCX, green) in hippocampal organoids generated from KOLF2-C1, KOLF2-1J and IMR90 iPSCs at d30 showing radial organisation in all iPSC lines, with centrally localised SOX2+ NSC layers, and a peripheral DCX+ immature neuron layer. Scale bar = 200 µm.

(**D**) Representative immunofluorescence images of dentate granule cells ($PROX1^{+}$, green) in hippocampal organoids generated from KOLF2-C1, KOLF2-1J and IMR90 iPSCs at d70. Scale bar = 50 µm.

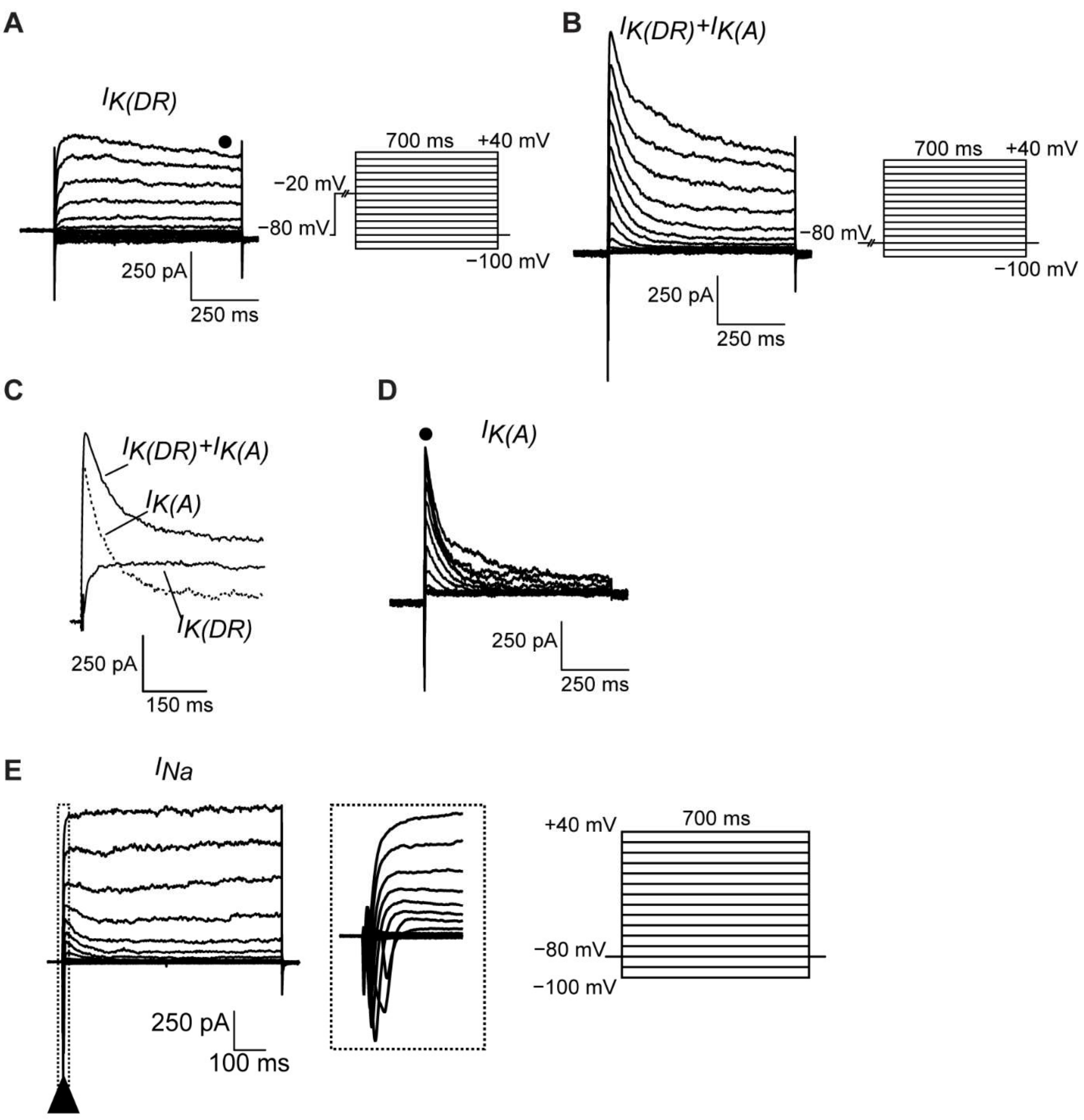


**Supplementary Figure 4. Voltage-clamp protocols and isolation of potassium and sodium currents in hippocampal organoid neurons. Related to Figure 4.**

(**A**) Voltage-clamp protocol used to isolate delayed rectifier potassium currents ($I_{K(DR)}$), consisting of a holding potential at −80 mV, a step to −20 mV, and subsequent depolarising steps from −100 mV to +40 mV in 10 mV increments.

(**B**) Voltage-clamp protocol used to measure total outward currents consisting of a holding potential at −80 mV and depolarising steps from −100 mV to +40 mV in 10 mV increments.

**(C)** Detail of current traces recorded from total outward currents and $I_{K(DR)}$, displaying how $I_{K(A)}$ is isolated by subtracting $I_{K(DR)}$ from total outward currents.

**(D)** Representative traces of transient A-type potassium currents ($I_{K(A)}$).

**(E)** Voltage-clamp protocol used to evoke sodium currents, involving a holding potential of −80 mV followed by depolarising steps from −100 to +40 mV in 10 mV increments. The amplitude of the transient inward current at the onset of each depolarisation step (triangle) was used to quantify *I(Na)*.

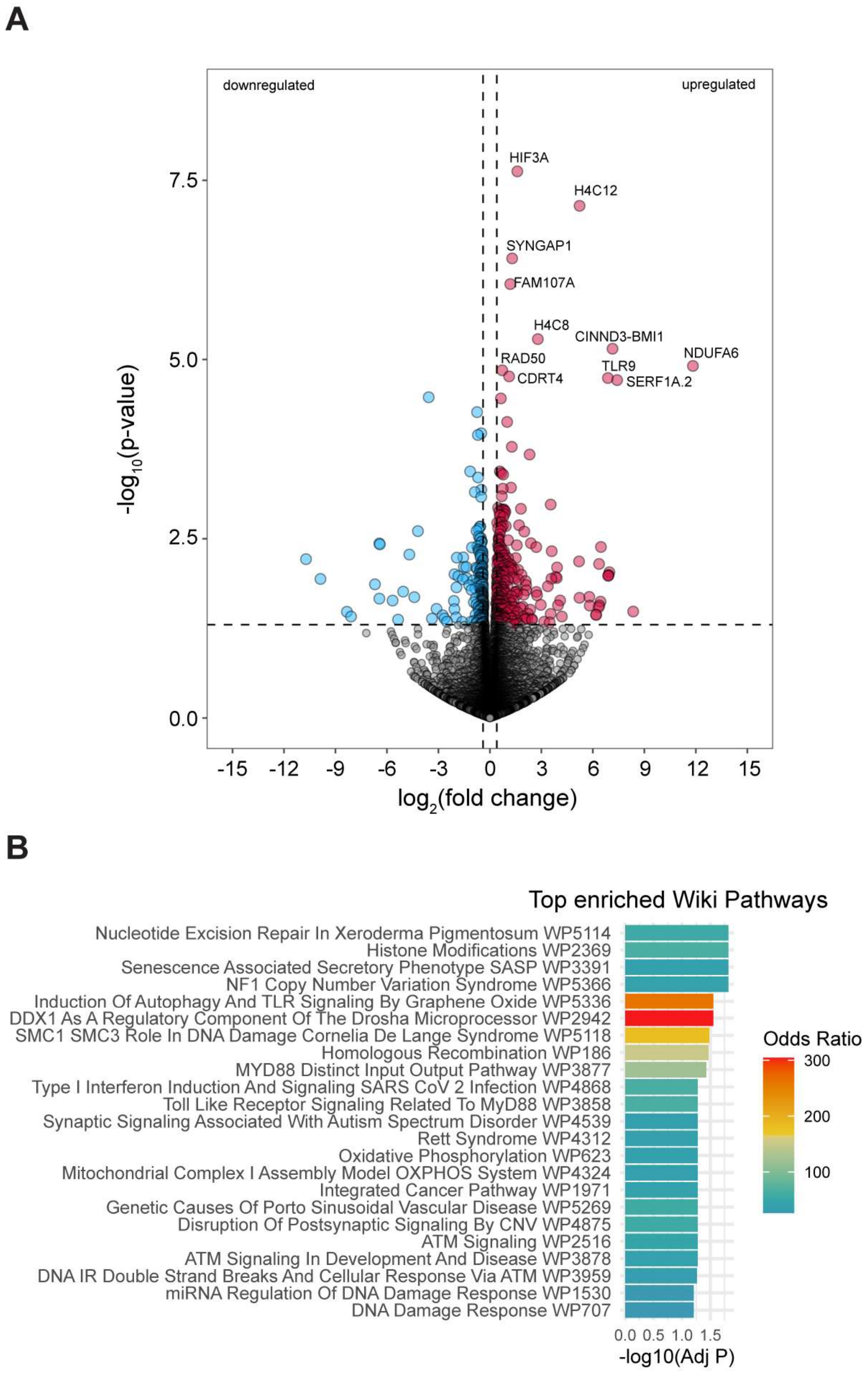
A
downregulated
upregulated
HIF3A
H4C12
SYNGAP1
FAM107A
H4C8
CINND3-BMI1
NDUFA6
RAD50
TLR9
CDRT4
SERF1A.2
-log10(p-value)
7.5
5.0
2.5
0.0
-15
-12
-9
-6
-3
0
3
6
9
12
15
log2(fold change)
B
Top enriched Wiki Pathways
Nucleotide Excision Repair In Xeroderma Pigmentosum WP5114
Histone Modifications WP2369
Senescence Associated Secretory Phenotype SASP WP3391
NF1 Copy Number Variation Syndrome WP5366
Induction Of Autophagy And TLR Signaling By Graphene Oxide WP5336
DDX1 As A Regulatory Component Of The Drosha Microprocessor WP2942
SMC1 SMC3 Role In DNA Damage Cornelia De Lange Syndrome WP5118
Homologous Recombination WP186
MYD88 Distinct Input Output Pathway WP3877
Type I Interferon Induction And Signaling SARS CoV 2 Infection WP4868
Toll Like Receptor Signaling Related To MyD88 WP3858
Synaptic Signaling Associated With Autism Spectrum Disorder WP4539
Rett Syndrome WP4312
Oxidative Phosphorylation WP623
Mitochondrial Complex I Assembly Model OXPHOS System WP4324
Integrated Cancer Pathway WP1971
Genetic Causes Of Porto Sinusoidal Vascular Disease WP5269
Disruption Of Postsynaptic Signaling By CNV WP4875
ATM Signaling WP2516
ATM Signaling In Development And Disease WP3878
DNA IR Double Strand Breaks And Cellular Response Via ATM WP3959
miRNA Regulation Of DNA Damage Response WP1530
DNA Damage Response WP707
0.0 0.5 1.0 1.5
-log10(Adj P)
Odds Ratio
300
200
100

**Supplementary Figure 5. Transcriptomic evidence of stress-associated pathways in dexamethasone-treated organoids. Related to Figure 7.**

(**A**) RNA-sequencing revealed transcriptional changes in dexamethasone-treated hippocampal organoids. Volcano plot showing the differentially expressed genes (DEGs). Among the DEGs were *SYNGAP1*, a known regulator of synaptic plasticity and also has roles in regulating the MAPK/ERK pathway, which is involved in regulating NSC proliferation, differentiation and survival. *H4C12* and *H4C8*, both histone H4 genes, were also differentially expressed and are implicated in chromatin regulation, DNA packaging, and DNA repair. Additionally, *RAD50*, a key component of the MRN complex involved in the repair of DNA double-strand breaks, was among the upregulated genes. Significantly upregulated genes (adjusted $p < 0.05$) are labelled; no significant downregulated genes detected. $n = 3$ organoids per group.

(**B**) DEGs were significantly enriched in pathways associated with cellular senescence and DNA damage responses. Specific upregulated pathways include senescence-associated secretory phenotype (WP3391), histone modifications (WP2369), homologous recombination (WP186), and DNA damage response (WP1530). $n = 3$ organoids per group.

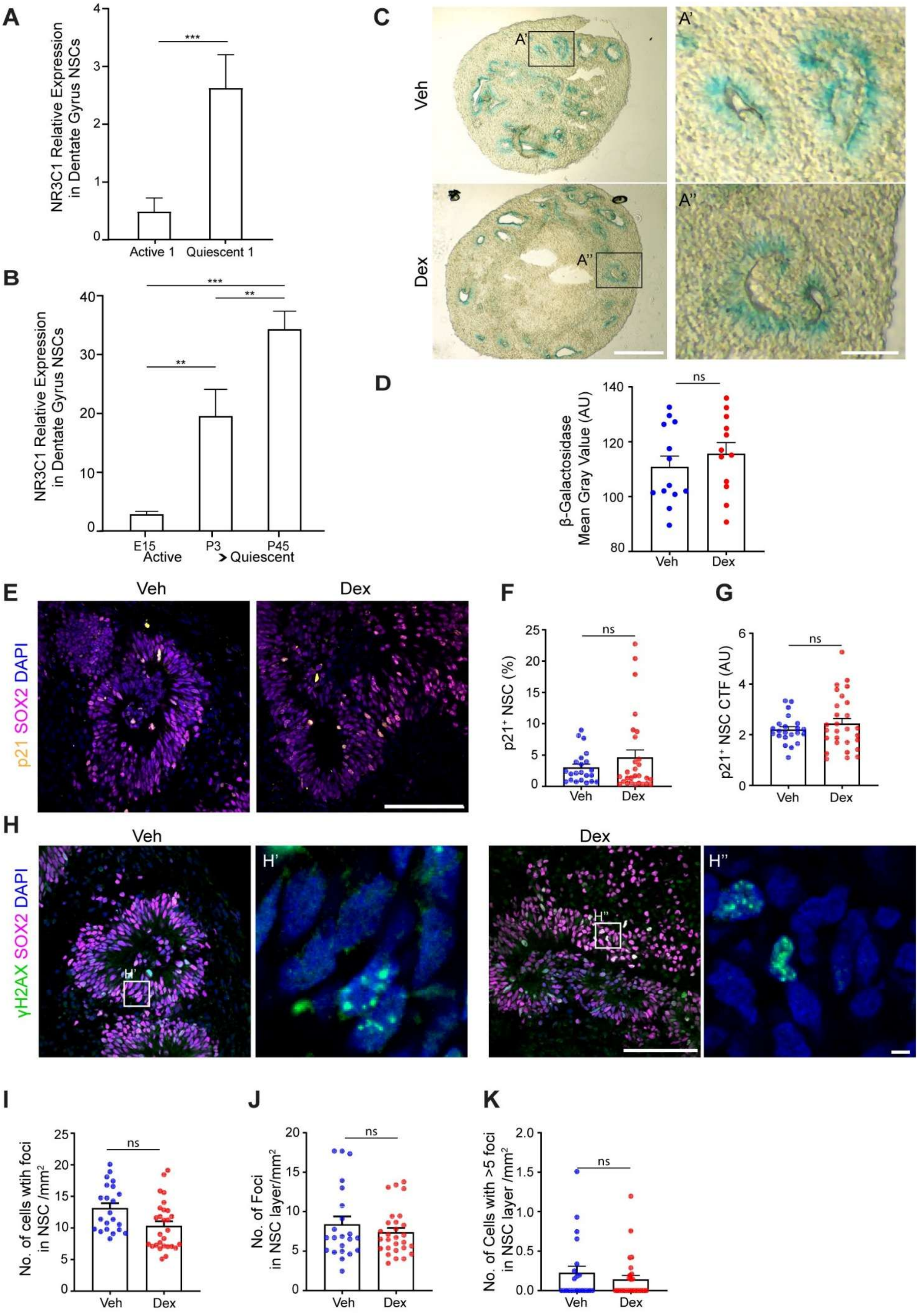
A
NR3C1 Relative Expression in Dentate Gyrus NSCs
***
Active 1
Quiescent 1
B
NR3C1 Relative Expression in Dentate Gyrus NSCs
***
**
**
E15
P3
P45
Active
Quiescent
C
Veh
Dex
A'
A''
D
β-Galactosidase Mean Gray Value (AU)
ns
Veh
Dex
E
Veh
Dex
p21 SOX2 DAPI
F
p21+ NSC (%)
ns
Veh
Dex
G
p21+ NSC CTF (AU)
ns
Veh
Dex
H
Veh
Dex
γH2AX SOX2 DAPI
H'
H''
I
No. of cells wtih foci in NSC /mm2
ns
Veh
Dex
J
No. of Foci in NSC layer/mm2
ns
Veh
Dex
K
No. of Cells with >5 foci in NSC layer /mm2
ns
Veh
Dex

**Supplementary Figure 6. Dexamethasone treatment does not induce senescence or DNA damage at the cellular level in hippocampal organoids. Related to Figure 7.**

**(A)** Relative expression of NR3C1 in active versus quiescent DG NSCs from Jimenez-Cyrus et al. Data are shown as mean ± SEM; n = 4–12 mice per timepoint. Significance was assessed using a Mann–Whitney test (***p = 0.0007).

**(B)** Relative expression of NR3C1 in DG NSCs at E15, P3, and P45 from Berg et al. Data are shown as mean ± SEM; n = 2–3 mice per timepoint. Significance was assessed using one-way ANOVA with Tukey's post hoc test (**p = 0.008, ***p = 0.0004).

**(C)** Representative brightfield images of β-galactosidase+ cells (blue), a canonical senescence marker, staining in vehicle- (veh) and dexamethasone- (dex) treated hippocampal organoids at d50. Positive cells for β-galactosidase were most abundant in the NSC layers with no observable difference in the spatial distribution of the β-galactosidase+ cells (inset). Scale bar = 1 mm.

**(D)** Quantification of β-galactosidase staining intensity, measured by mean gray value, revealed no significant differences were between veh- and dex-treated organoids. Data are presented as mean + SEM. n = 12 – 13 images from 3 hippocampal organoids per treatment group at d50. Statistical significance determined by unpaired t-test. p = 0.4097.

**(E)** Representative immunofluorescence images of p21+ (orange) in NSCs (SOX2, magenta) in veh-and dex-treated hippocampal organoids at d50. p21 is a multifunctional protein with broad regulatory roles in a range of cellular processes, including cell cycle progression, apoptosis, differentiation, cell migration, cytoskeletal organisation, transcriptional regulation, DNA repair, and the initiation of cellular senescence. Scale bar = 100 μm.

**(F-G)** Treatment with dexamethasone does not alter the total proportion of p21+ NSCs (p = 0.5675) (**F**) or the intensity of the p21+ signal in NSCs, measured by mean gray value (p = 0.2722) (**G**). Data are presented median ± IQR. n = 22 – 28 images from 3 hippocampal organoids per treatment group at d50. Statistical significance determined by Mann-Whitney test.

**(H)** Representative immunofluorescence images of double strand DNA breaks (γH2AX, green) in neural stem cells (SOX2, magenta) in veh- and dex-treated hippocampal organoids. Magnified views of γH2AX immunoreactivity highlighting individual foci are shown in H′ and H″. γH2AX, which is generated by phosphorylation of the histone variant H2AX at serine 139, is well-established as an early and sensitive marker of DNA double-strand breaks and is commonly used to monitor the initiation and resolution of DNA damage. Upon DNA damage, γH2AX forms discrete nuclear foci at the sites of breaks, which recruit and anchor DNA repair proteins to facilitate the damage response. Scale bar = 100 μm.

**(I-K)**. Treatment with dexamethasone does not increase the number of NSCs with foci (p = 0.0105) (**I**), the number of foci per NSC layer (p = 0.7346) **(J**) or the number of cells per NSC layer with 5 or more foci (p = 0.5711) (**K**). Data are presented as median ± IQR. n = 22 – 28 images from 3 hippocampal organoids per treatment group. Statistical significance determined by Mann-Whitney test.
